# Global Signal Regression Strengthens Association between Resting-State Functional Connectivity and Behavior

**DOI:** 10.1101/548644

**Authors:** Jingwei Li, Ru Kong, Raphael Liegeois, Csaba Orban, Yanrui Tan, Nanbo Sun, Avram J. Holmes, Mert R. Sabuncu, Tian Ge, B.T. Thomas Yeo

## Abstract

Global signal regression (GSR) is one of the most debated preprocessing strategies for resting-state functional MRI. GSR effectively removes global artifacts driven by motion and respiration, but also discards globally distributed neural information and introduces negative correlations between certain brain regions. The vast majority of previous studies have focused on the effectiveness of GSR in removing imaging artifacts, as well as its potential biases. Given the growing interest in functional connectivity fingerprinting, here we considered the utilitarian question of whether GSR strengthens or weakens associations between resting-state functional connectivity (RSFC) and multiple behavioral measures across cognition, personality and emotion.

By applying the variance component model to the Brain Genomics Superstruct Project (GSP), we found that behavioral variance explained by whole-brain RSFC increased by an average of 47% across 23 behavioral measures after GSR. In the Human Connectome Project (HCP), we found that behavioral variance explained by whole-brain RSFC increased by an average of 40% across 58 behavioral measures, when GSR was applied after ICA-FIX de-noising. To ensure generalizability, we repeated our analyses using kernel regression. GSR improved behavioral prediction accuracies by an average of 64% and 12% in the GSP and HCP datasets respectively. Importantly, the results were consistent across methods. A behavioral measure with greater RSFC-explained variance (using the variance component model) also exhibited greater prediction accuracy (using kernel regression). A behavioral measure with greater improvement in behavioral variance explained after GSR (using the variance component model) also enjoyed greater improvement in prediction accuracy after GSR (using kernel regression). Furthermore, GSR appeared to benefit task performance measures more than self-reported measures.

Since GSR was more effective at removing motion-related and respiratory-related artifacts, GSR-related increases in variance explained and prediction accuracies were unlikely the result of motion-related or respiratory-related artifacts. However, it is worth emphasizing that the current study focused on whole-brain RSFC, so it remains unclear whether GSR improves RSFC-behavioral associations for specific connections or networks. Overall, our results suggest that at least in the case for young healthy adults, GSR strengthens the associations between RSFC and most (although not all) behavioral measures. Code for the variance component model and ridge regression can be found here: https://github.com/ThomasYeoLab/CBIG/tree/master/stable_projects/preprocessing/Li2019_GSR.

**Highlights:**

1. Global signal regression improves RSFC-behavior associations
2. Global signal regression improves RSFC-based behavioral prediction accuracies
3. Improvements replicated across two large-scale datasets and methods
4. Task-performance measures enjoyed greater improvements than self-reported ones
5. GSR beneficial even after ICA-FIX

## 1. Introduction

Resting-state functional connectivity (RSFC) is a powerful tool for measuring the synchronization of fMRI signals between brain regions, while participants are lying at rest without any “extrinsic” task (Biswal et al., 1995; Fox and Raichle 2007; Buckner et al., 2013). RSFC has been widely used to delineate large-scale brain networks and explore human brain organization (Fox et al., 2005; Seeley et al., 2007; Smith et al., 2009; Bertolero et al., 2015; Eickhoff et al., 2018). However, fMRI is contaminated by various noise sources, which have been shown to be particularly problematic for resting-state functional MRI (rs-fMRI) studies (Power et al., 2012; Satterthwaite et al., 2012; Van Dijk et al., 2012; Yan et al., 2013). Consequently, there has been significant research on rs-fMRI denoising (Chai et al., 2012; Hallquist et al., 2013; Jo et al., 2013; Patel et al., 2014; Bright et al., 2017), but there is still no consensus approach for rs-fMRI preprocessing (Murphy et al., 2013; Geerligs et al., 2017; Murphy and Fox, 2017).

One of the most contentious rs-fMRI preprocessing steps is global signal regression (GSR; Uddin 2017; Power et al., 2017a). Although the exact implementation varies across studies (Liu et al., 2017), GSR typically involves the regression of whole brain (or gray matter or cortical) fMRI signal from every brain voxel (Power et al., 2014; Burgess et al., 2016; Power et al., 2018). The global signal is associated with head motion, respiration and cardiac rhythms (Birn et al., 2006; Power et al., 2014; 2017; Liu et al., 2017). Consequently, GSR is highly effective at removing global artifacts arising from motion and other physiological sources. More specifically, studies have shown that GSR reduces the correlation magnitude between quality control metrics and RSFC (Satterthwaite et al., 2013; Power et al., 2014; Burgess et al., 2016; Ciric et al., 2016; Parkes et al., 2018), as well as removes prominent increases and/or decreases in signal intensities lasting many TRs (Power et al., 2014; Byrge et al., 2017; Glasser et al., 2018). GSR also improves neuronal-hemodynamic correspondence (Keller et al., 2013).

However, GSR has been criticized to introduce negative correlations, so the signs of the resulting correlations might be hard to interpret, i.e., negative correlations after GSR might be a mathematical consequence of GSR, rather than actual inhibitory interactions between brain regions (Murphy et al., 2009; Weissenbacher et al., 2009). GSR might also distort group differences (Saad et al., 2012). For example, the global signal differs between certain populations, so GSR might remove RSFC differences between diseased populations and healthy controls (Yang et al., 2014; but see Parkes et al., 2018). Furthermore, because the global signal itself contains neuronal information (Schölvinck et al., 2010; Matsui et al., 2016; Wen and Liu, 2016) and is associated with vigilance and arousal (Wong et al., 2013, 2016; Yeo et al., 2015b; Chang et al., 2016), some have argued against removing such information (Cole et al., 2010; Ben Simon et al., 2017; Glasser et al., 2018). Finally, although GSR reduces the correlation magnitude between quality control metrics and RSFC, this reduction is distance dependent, which might potentially introduce biases in certain analyses (Power et al., 2014; Parkes et al., 2018).

A different, but potentially useful, perspective is that GSR simply changes the “reference” signal, and hence what is being measured. More specifically, analyses without GSR evaluates total fMRI fluctuation (global signal fluctuations plus fluctuations relative to the global signal), while analyses with GSR examines fMRI fluctuations relative to the global signal (Yeo et al., 2015b). Therefore, whether GSR should be performed or not depends on the scientific or clinical question being asked. Indeed, when using electroencephalogram (EEG) to detect seizures, the “best” reference electrode(s) is the one that makes the epileptiform discharge most visible to the clinician (Murphy and Fox, 2017). Similarly, removing the global gene expression pattern might also be useful for genetic analyses (Krienen et al., 2016; Anderson et al., 2018; Filbin et al., 2018). Therefore, under this perspective, whether GSR should be performed depends on its utility in the problem being studied.

Given significant interest in the relationship between behavior and functional brain architecture revealed by RSFC (Fox et al., 2012; Mueller et al., 2013; Smith et al., 2015; Finn et al., 2015; Rosenberg et al., 2016; Dubois et al., 2016; Bertolero et al., 2018; Kong et al., 2018), here we explored the utilitarian question of whether GSR strengthens or weakens the association between RSFC and behavior in young healthy adults. In the literature, results on whether GSR strengthens or weakens RSFC-behavioral association are mixed. While some studies have suggested that GSR leads to stronger relationships between RSFC and behavior (Hampson et al., 2010; Kruschwitz et al., 2015; Yeo et al., 2015b), others have suggested that GSR weakens relationship between RSFC and behavior, especially when comparing healthy controls and diseased populations (Gotts et al., 2013; Yang et al., 2014; but see Parkes et al., 2018).

In this work, we utilized the variance component model (Yang et al., 2011; Sabuncu et al., 2016) to quantify the association between behavior and RSFC with and without GSR. The variance component model has been widely used to estimate the variance explained by genome-wide genetic variants for a complex trait (Yang et al., 2011), and was recently applied to associating neuroimaging data with behavioral measurements (Sabuncu et al., 2016). To ensure our conclusions were robust to the choice of analysis strategy, we repeated the analyses using kernel ridge regression for RSFC-based behavioral prediction (He et al., 2018). Kernel ridge regression provides a fast and effective way for predicting behavioral phenotype in individual subjects using rs-fMRI (He et al., 2018; Kong et al., 2018). Here, we applied both approaches (variance component model and kernel ridge regression) to two large-scale datasets of young healthy adults: the Brain Genomics Superstruct Project (GSP; Holmes et al., 2015) and the Human Connectome Project (HCP; Van Essen et al., 2013). Because the two datasets were collected using different scanner type (e.g., Trio versus Skyra) and acquisition sequence (e.g., multiband versus non-multiband), this allows us to test whether the effects are robust across acquisition parameters. In the case of the HCP dataset, it has previously been denoised using ICA-FIX (Salimi-Khorshidi et al., 2014; Griffanti et al., 2014). This allows us to test whether GSR was useful above and beyond ICA-FIX.

## 2. Methods and Materials

### 2.1 Datasets and preprocessing

Two publicly available datasets were examined: Brain Genomics Superstruct Project (GSP; Holmes et al., 2015) and Human Connectome Project (HCP) S1200 release (Van Essen et al., 2013). Both datasets contained structural MRI, rs-fMRI, and multiple behavioral measures for each subject.

All participants in the GSP dataset were healthy and young (ages 18-35). MR images were collected on matched 3T Tim Trio scanners (Siemens Healthcare, Erlangen, Germany) at Harvard University and Massachusetts General Hospital using the vendor-supplied 12-channel phased-array head coil. The T1-weighted structural images were 1.2mm isotropic. Rs-fMRI data were 3mm isotropic with TR = 3000ms. Each rs-fMRI scan has 124 frames. Most participants (N = 1490) had one session of rs-fMRI data, while a small number of participants (N = 69) had two sessions of data collected less than 6 months apart. Each session included one or two rs-fMRI runs. The behavioral data were collected from computer-based cognitive tasks and personality assessment. Behavioral measurements of non-compliant participants were removed (Holmes et al., 2015). Further details can be found elsewhere (Holmes et al., 2015).

HCP participants (N = 1094) were drawn from a population of twins and siblings. Participants were healthy and young (ages 22-37). All imaging data were acquired on a customized Siemens 3T Skyra at Washington University (St Louis) using a multi-band sequence. The structural images were 0.7mm isotropic. The rs-fMRI data were 2mm isotropic with TR = 0.72s. Two sessions of rs-fMRI data were collected in consecutive days for each subject, and each session consisted of one or two runs. The length of each rs-fMRI scan was 14.4 min (1200 frames). Details of the data collection can be found elsewhere (Van Essen et al., 2012; Smith et al., 2013). Details about behavioral measures can be found in HCP S1200 Data Dictionary and Barch et al. (2013).

#### 2.1.1 GSP preprocessing and behavioral data

The T1 images of the GSP dataset were previously processed (Holmes et al., 2015) using FreeSurfer 4.5.0 recon-all procedure (http://surfer.nmr.mgh.harvard.edu; Dale et al., 1999; Ségonne et al., 2004, 2007; Fischl et al., 1999a, 1999b, 2001). FreeSurfer provides automatic algorithms for cortical reconstruction and volumetric segmentation from individual subjects’ T1 images (Dale et al., 1999; Fischl et al., 2002; Segonnne et al., 2007). Each subject’s cortical surface mesh was registered to a common spherical coordinate system (Fischl et al., 1999a, 1999b).

Two preprocessing pipelines were applied to rs-fMRI data: GSP-Baseline and GSP-Baseline+GSR. The pipelines consisted of a combination of FreeSurfer 5.3.0 (Fischl et al., 2012), FSL 5.0.8 (Jenkinson et al., 2012; Smith et al., 2004) and in-house Matlab functions. Both pipelines underwent the following steps: (1) removal of the first four frames, (2) slice time correction with FSL package, (3) motion correction using rigid body translation and rotation using FSL, together with outlier detection (see below), (4) alignment with structural image using boundary-based registration (Greve and Fischl, 2009) provided by FsFast (http://surfer.nmr.mgh.harvard.edu/fswiki/FsFast), (5) nuisance regression (details below), (6) interpolation of censored frames with Lomb-Scargle periodogram (Power et al., 2014), (7) band-pass filtering (0.009 Hz ≤ f ≤ 0.08 Hz), (8) projection onto the FreeSurfer fsaverage6 surface space and (9) smoothing by a 6mm full-width half-maximum kernel (FWHM). The only difference between the two pipelines (GSP-Baseline and GSP-Baseline+GSR) occurred in the nuisance regression step. Details of outlier detection and nuisance regression are elaborated below.

Framewise displacement (FD; Jenkinson et al., 2002) and root-mean-square of voxel-wise differentiated signal (DVARS) (Power et al., 2012) were estimated using fsl_motion_outliers. Volumes with FD > 0.2mm or DVARS > 50 were marked as outliers (censored frames). One frame before and two frames after these volumes were flagged as censored frames. Uncensored segments of data lasting fewer than five contiguous volumes were also labeled as censored frames (Gordon et al., 2016). BOLD runs with more than half of the volumes labeled as censored frames were removed.

Linear regression of multiple nuisance regressors was applied. In the GSP-Baseline pipeline, nuisance regressors consisted of: (1) a vector of ones and linear trend, (2) six motion correction parameters, (3) averaged white matter signal, (4) averaged ventricular signal, and (5) temporal derivatives of (2) - (4). The white matter mask was obtained from FreeSurfer’s segmentation of the anatomical T1 image, followed by three rounds of erosion before projection to the subject’s native rs-fMRI space. The ventricle mask was obtained in a similar fashion, but only with one round of erosion. If there were less than 100 voxels (in anatomical space) after erosion, then no erosion was performed. In the GSP-Baseline+GSR pipeline, in addition to the previously mentioned regressors, mean whole brain signal and its temporal derivative were also removed. Across both pipelines, censored frames were ignored when computing the regression coefficients (Power et al., 2014).

Functional connectivity (FC) was computed among 419 ROIs using Pearson’s correlation. The 419 ROIs consisted of 400 cortical parcels (Schaefer et al., 2017; https://github.com/ThomasYeoLab/CBIG/tree/master/stable_projects/brain_parcellation/Schaefer2018_LocalGlobal) and 19 subcortical ROIs in subject-specific volumetric space defined by FreeSurfer (Fischl et al., 2002). The 19 subcortical ROIs corresponded to the cerebellar gray matter, thalamus, caudate, putamen, pallidum, hippocampus, accumbens, amygdala, ventral diencephalon, and brain stem. Censored frames were ignored when computing the correlations. For each subject, the correlation matrix was computed for each run, Fisher’s z-transformed, and then averaged across runs and sessions, yielding one final 419 × 419 RSFC matrix for each subject.

A total of 1551 subjects survived the fMRI preprocessing quality control (e.g., motion censoring). 23 behavioral phenotypes, including cognitive and personality measures, were considered (Table S1). 689 participants were further excluded because they did not have all 23 behavioral measurements, yielding a final set of 862 participants.

#### 2.1.2 HCP preprocessing and behavioral data

Details of the HCP preprocessing can be found elsewhere (HCP S1200 manual; Van Essen et al. 2012; Glasser et al. 2013; Smith et al. 2013). Of particular importance is that both cortical and subcortical data were denoised with ICA-FIX (Salimi-Khorshidi et al., 2014; Griffanti et al., 2014) and saved in the CIFTI grayordinate format. The surface (fs_LR) data were aligned with MSM-All (Robinson et al., 2013).

Recent studies have suggested that ICA-FIX does not fully eliminate global motion-related and respiratory-related artifacts (Burgess et al., 2016; Siegel et al., 2017), so further censoring was performed. Volumes with FD > 0.2mm or DVARS > 75 were marked as outliers (censored frames). One frame before and two frames after these volumes were flagged as censored frames. Uncensored segments of data lasting fewer than five contiguous volumes were also labeled as censored frames. BOLD runs with more than half of frames flagged as censored were removed. We refer to this data as being processed by the HCP-Baseline pipeline.

To examine the effects of global signal regression, we additionally regressed the global signal obtained by averaging across all cortical vertices^1^ and its temporal derivative (Power et al., 2018). Like before, censored frames were ignored when regression coefficients were computed. We refer to this processing pipeline as HCP-Baseline+GSR.

FC was computed among 419 ROIs using Pearson’s correlation. The 400 cortical ROIs were the same as before, but in fs_LR space (Schaefer et al., 2017; https://github.com/ThomasYeoLab/CBIG/tree/master/stable_projects/brain_parcellation/Schaefer2018_LocalGlobal). The 19 subcortical ROIs were defined based on grayordinate data structure. Pearson’s correlations were calculated only with the uncensored frames. For each subject, the correlation matrix was computed for each run, Fisher’s z-transformed, and then averaged across runs and sessions, yielding one final 419 × 419 RSFC matrix for each subject.

58 behavioral measures across cognition, personality and emotion were selected (Table S2; Kong et al., 2018). Of the 1029 subjects who survived the motion censoring, an additional 76 participants were excluded due to missing data for the behavioral measures, resulting in 953 subjects.

#### 2.1.3 QC-FC correlations for preprocessed rs-fMRI

We first replicated previous literature showing that GSR reduces imaging-related artifacts, while increasing distance-dependent biases. In addition to the individual subject QC plots pioneered by Power and colleagues (Power et al., 2014; Power, 2017), we also computed QC-FC correlation metrics widely used in the literature (Satterthewaite et al., 2012; Power et al., 2014; Burgess et al., 2016; Ciric et al., 2017). For each ROI pair, the QC-FC correlation was defined as the across-subject correlation between the functional connectivity of the ROI pair and the number of censored frames. The number of censored frames was chosen as the QC measure because it reflected both FD and DVARS, thus taking into account both head motion and other noise sources causing frame-to-frame signal intensity fluctuations (e.g. respiration).

### 2.2 Variance component model

The association between functional connectivity and a behavioral phenotype (e.g., fluid intelligence) was captured by a variance component model (Yang et al., 2011; Sabuncu et al., 2016):

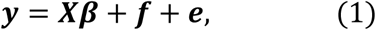

where ***y*** is an *N* × 1 vector of behavioral phenotype and *N* is the number of subjects in the dataset. ***N*** is an *N* × *P* nuisance covariate matrix. ***β*** denote the *P* regression coefficients. Age, sex, and motion (FD and DVARS) were included as covariates. Subject-specific effect ***f*** follows a multivariate normal distribution 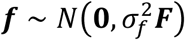, where 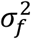 is a scaling constant (estimated from data) and ***F*** is the *N* × *N* functional connectivity similarity matrix among subjects. More specifically, the *i*-th row and *j*-th column of ***F*** corresponds to the functional connectivity similarity between the *i*-th and *j*-th subjects. In the variance component model, the diagonal entries of the matrix ***F*** are assumed to be one (Yang et al., 2011; Sabuncu et al., 2016). Inspired by previous RSFC fingerprinting studies that identified individuals based on the Pearson’s correlation between their RSFC matrices (Finn et al., 2015), here we defined the functional connectivity similarity between the *i*-th and *j*-th subjects to be the Pearson’s correlation between their 419 × 419 RSFC matrices (considering only lower triangular entries since the matrices are symmetric).

Overall, the intuition behind the variance component model is that subjects *i* and *j* have more similar behavior if their RSFC matrices are more similar. The degree in which this assumption is true is reflected by the scaling constant 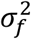. The error behind this assumption is captured by the noise component ***e***, which is assumed to be i.i.d. Gaussian distributed for each subject, i.e., 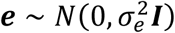, where ***I*** is an identity matrix. In other words, if 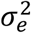 is large relative to 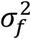, this would imply that individual differences in a behavioral phenotype are not well captured by individual differences in RSFC. This motivates the following definition of the variance of ***y*** explained by RSFC (Yang et al., 2011; Sabuncu et al., 2016):

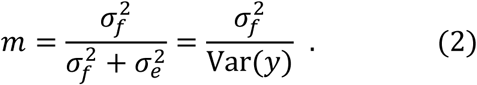

A larger *m* implies that a larger portion of the individual differences in the behavioral phenotype can be explained by individual differences in RSFC. In practice, 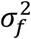 and 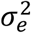 were iteratively estimated using the restricted maximum likelihood (ReML) algorithm (Sabuncu et al., 2016).

We applied the variance component model to all 23 behavioral measures, age and sex in the GSP dataset. All participants (N = 862) were utilized. We also applied the variance component model to all 58 behavioral measures, age and sex in the HCP dataset. Because dealing with both RSFC and family structure within the variance component model is tricky, we followed the strategy of choosing one subject from each family (Sabuncu et al., 2016), resulting in 419 unrelated subjects. When analyzing age (or sex), age (or sex) was not included as a nuisance covariate.

Since the variance component model assumes the behavioral scores are Gaussian distributed, each behavioral phenotype was quantile normalized before the variance component model was fitted (Elliott et al., 2018). To perform quantile normalization for a particular behavioral phenotype, the scores of all participants were sorted from low to high. The scores were then replaced by samples from a standard Gaussian distribution (also sorted from low to high). Thus, the resulting behavioral distribution follows a Gaussian distribution. We note that not performing quantile normalization yielded very similar results.

A jackknife procedure was utilized to quantify the uncertainty of the variance estimates (Ge et al., 2016). For each behavioral phenotype, half the participants were randomly removed. The variance component model was fitted for the remaining participants. This jackknife procedure was repeated 1000 times, resulting in 1000 jackknife estimates of the explained behavioral variance. Because half the participants were deleted in each jackknife sample, the sample variance of the 1000 jackknife estimates can be used as an estimate of the uncertainty of the explained behavioral variance (Shao, 1989; Shao and Tu, 2012).

### 2.3 Kernel ridge regression

To ensure our conclusions are robust to the choice of analysis strategy, we utilized kernel ridge regression (Murphy, 2012) to compare RSFC-based behavioral prediction accuracies with and without GSR. Suppose *y*_*s*_ and *y*_*i*_ denote the behavioral measure (e.g. fluid intelligence) of test subject *s* and training subject *i* respectively. Let *c*_*s*_ and *c*_*i*_ denote the vectorized RSFC (lower triangular entries of the RSFC matrices) of test subject *s* and training subject *i* respectively. Then, roughly speaking, kernel regression would predict *y*_*s*_ as the weighted average of the behavioral measures of all training subjects, i.e., 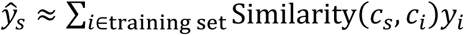. Here, we set Similarity(*c*_*s*_, *c*_*i*_) to be the Pearson’s correlation between the vectorized RSFC of the test subject and the *i*-th training subject. Therefore, successful prediction would indicate that subjects with more similar RSFC have similar behavioral scores. To reduce overfitting, an *l*_2_-regularization term was included, i.e., kernel ridge regression was utilized. More details can be found in Appendix I.

We applied kernel ridge regression to predict all 23 behavioral measures, age and sex in the GSP dataset. All participants (N = 862) were utilized. We also applied kernel ridge regression to predict all 58 behavioral measures, age and sex in the HCP dataset. For both datasets, we performed 20-fold cross-validation for each behavioral (or demographic) measure. For each test fold, the kernel ridge regression parameters were estimated from the remaining 19 training folds. 20-fold cross-validation was in turn performed on the 19 training folds with different *l*_2_-regularization parameter *λ*, in order to select the best *λ*. The estimated parameters from the training folds were then used to predict behavior of the subjects in the test fold.

For both datasets, all participants were utilized. Family structure within the HCP dataset was taken into account by ensuring participants from the same family were not split across folds. Prediction accuracies were measured by the Pearson’s correlation between the raw behavioral score and the predicted score across all subjects in a test fold. Since there were 20 test folds, there were 20 correlation accuracies for each behavioral measure. These 20 accuracies were then averaged across the test folds. Because a single 20-fold cross-validation might be sensitive to the particular split of the data into folds (Varoquaux et al., 2017), the above 20-fold cross-validation was repeated 20 times.

Because certain behavioral measures are known to correlate with motion (Siegel et al., 2016), we regressed age, sex, FD and DVARS from each behavioral measure before kernel ridge regression. The regression coefficients were calculated from training set and applied to the test set. When predicting age (or sex), age (or sex) was not included in the regressors.

Finally, we note that linear ridge regression was also considered. The results were consistent with all conclusions in this paper. Because of the current length of this manuscript and because kernel ridge regression accuracies were better than linear ridge regression, we decided to focus on kernel ridge regression for the remainder of this paper.

### 2.4 Consistencies and heterogeneities across methods and behavioral measures

#### 2.4.1 Consistency of results across methods

We first explored whether the results were consistent between variance component model and kernel ridge regression. For each dataset, we correlated the RSFC-explained variances and prediction accuracies across behavioral measures. In other words, we wanted to explore whether a behavioral measure better explained by RSFC (using the variance component model) was also more easily predicted by kernel ridge regression.

Second, for each dataset, we correlated changes in explained variances (with and without GSR) and changes in prediction accuracies (with and without GSR) across behavioral measures. In other words, we wanted to explore whether a behavioral measure with a larger improvement in RSFC-explained variance after GSR (using the variance component model) will also enjoy greater improvement in prediction accuracy after GSR.

#### 2.4.2 Relationship with baseline associations and prediction accuracies

To explore which behavioral measures benefited most from GSR, we hypothesized that GSR-related changes might be related to baseline RSFC-behavioral associations or prediction accuracies. For example, we could imagine that behavioral measures that were more poorly predicted using the baseline preprocessing strategy might benefit more from GSR and vice versa. Thus, for each dataset, we correlated changes in explained variances (with and without GSR) with the explained variances of the baseline preprocessing pipeline across behavioral measures. The analysis was repeated with prediction accuracies.

#### 2.4.3 Task-performance and self-reported measures

To further explore which behavioral measures benefited most from GSR, a second hypothesis was that GSR-related improvements might be related to whether a behavioral measure involved a cognitive task or was a self-reported measure. Thus, the 58 HCP behavioral measures were grouped into 27 “task-performance” measures, 24 “self-reported” measures and 7 unclassified measures. The task-performance measures were defined as the tasks that assess cognitive performance satisfying two criteria. First, there were some cognitive processes involved when the participants performed the task. Second, the task was performance-based, i.e. the behavioral scores reflected whether the participants performed the task well or not. On the other hand, self-reported measures were those in which participants provided answers to the surveys themselves. The remaining measures were considered as unclassified. For example, the delay discounting scores were collected based on a task, but a subject with higher score does not mean “better” performance, thus delay discounting was considered unclassified. Similarly, the odor identification measure was performance-based, but there was little cognitive processing in odor identification, so the measure was also considered unclassified. For the HCP dataset, we then explored whether GSR-related improvements (in explained variance and prediction accuracies) differed between task-performance and self-reported measures. In the case of the GSP dataset, there were too few task-performance measures to perform a similar analysis.

### 2.5 Further motion analyses

In this section, we discuss further motion controls and analyses. First, in the original analyses (Section 2.2), FD and DVARS were included as nuisance covariates in the variance component model. We explored the effects of not including FD and DVARS as covariates. One hypothesis is that not including FD and DVARS as covariates might increase the explained variance since some behavioral measures are correlated with motion (Siegel et al., 2016), which might in turn correlate with residual motion artifacts in the rs-fMRI data.

Second, we computed correlations between all behavioral measures and motion (FD or DVARS). We then compared the magnitude of the correlations with the explained behavioral variance (variance component model) and the behavioral prediction accuracies (kernel ridge regression), in order to show that the explained variance and prediction accuracies cannot be entirely explained by motion.

### 2.6 Data and code availability

Code for this work is freely available at the GitHub repository maintained by the Computational Brain Imaging Group (https://github.com/ThomasYeoLab/CBIG). The preprocessing pipeline for the GSP dataset can be found here (https://github.com/ThomasYeoLab/CBIG/tree/master/stable_projects/preprocessing/CBIG_fMRI_Preproc2016). The Schaefer parcellation can be found here (https://github.com/ThomasYeoLab/CBIG/tree/master/stable_projects/brain_parcellation/Schaefer2018_LocalGlobal). The code for the variance component model and kernel ridge regression can be found here (https://github.com/ThomasYeoLab/CBIG/tree/master/stable_projects/preprocessing/Li2019_GSR). Both the GSP (http://neuroinformatics.harvard.edu/gsp/) and HCP (https://www.humanconnectome.org/) datasets are publicly available.

## 3. Results

### 3.1 Overview

In Section 3.2, we replicated results in the literature, showing that GSR introduces negative correlations, while reducing imaging artifacts. This is followed by variance component results and behavioral prediction results in Sections 3.3 and 3.4 respectively. In Section 3.5, we explored consistencies and heterogeneities across approaches and behavioral measures. Finally, motion-related effects were investigated in Section 3.6.

### 3.2 GSR effects on RSFC and fMRI timeseries

#### 3.2.1 Consistent with the literature, GSR reduces RSFC strength

Figure 1A shows the 400-area cortical parcellation (Schaefer et al., 2017). The colors corresponded to the 17 large-scale networks from Yeo et al. (2011). Figure 1B shows the 19 subcortical ROIs (Fischl et al., 2002). Figures 1C and 1D show the functional connectivity among the 419 ROIs averaged across participants with and without GSR in the GSP dataset. The results for the HCP dataset are shown in Figure S1.

**Figure 1.**
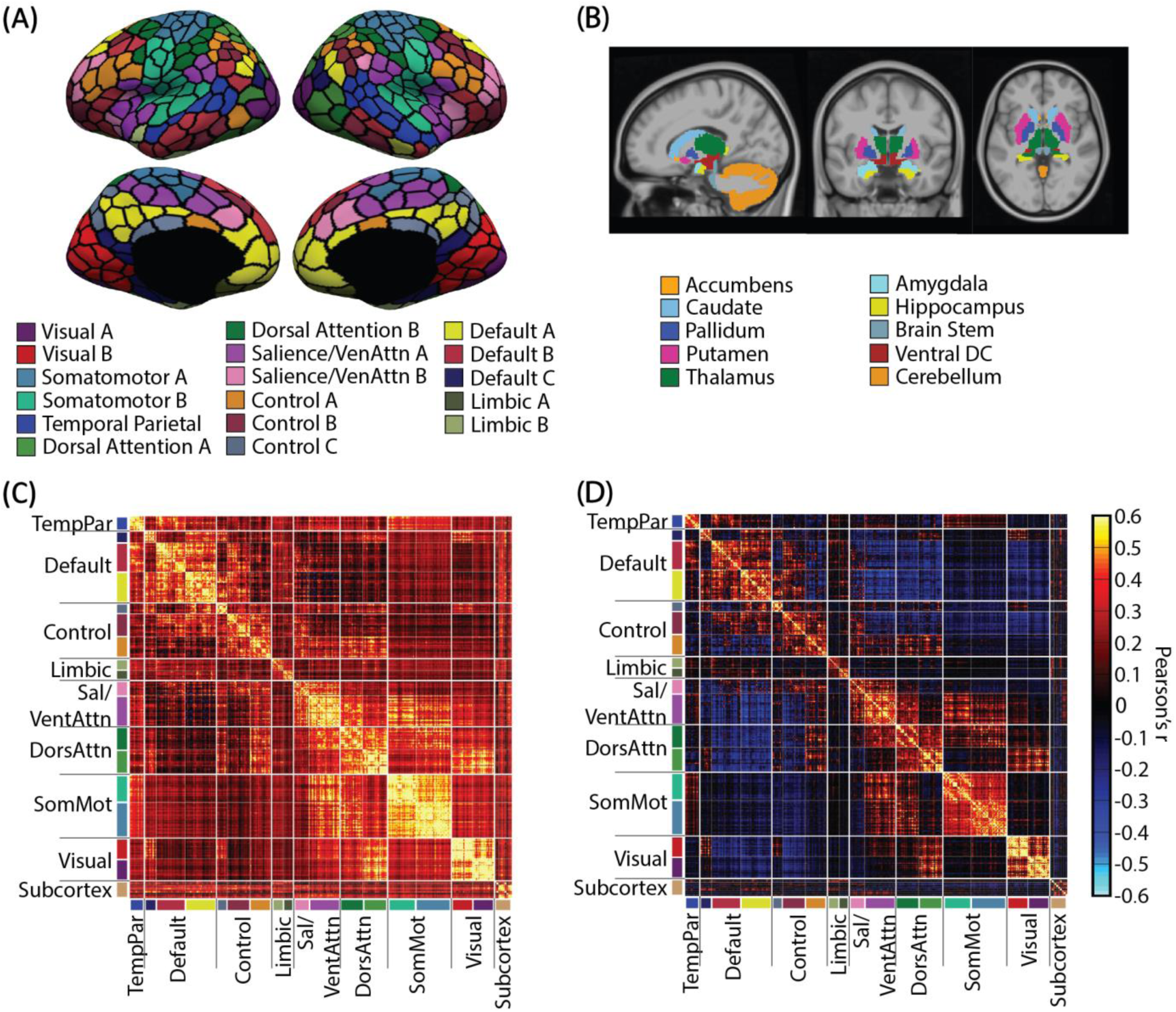
GSR results in a negative “shift” in RSFC in the GSP dataset. (A) 400-area cortical parcellation (Schaefer et al., 2017). Parcel colors correspond to 17 large-scale networks (Yeo et al., 2011). (B) 19 subcortical ROIs (Fischl et al., 2002). (C) RSFC matrix among the 419 ROIs using baseline processing without GSR. (D) RSFC matrix among the 419 ROIs using baseline processing with GSR. For visualization, the 419 ROIs are ordered according to the 17 networks in (A) and subcortical structures listed in (B). These 17 networks are in turn divided into eight groups (TempPar, Default, Control, Limbic, Salience/Ventral Attention, Dorsal Attention, Somatomotor and Visual), roughly corresponding to major networks discussed in the literature. These eight groups and subcortical structures are separated by thick while lines. Consistent with the literature, GSR introduces a negative shift in the RSFC.

In both GSP and HCP datasets, the RSFC patterns were highly similar with and without GSR. Correlations between the 419 × 419 RSFC matrices (excluding diagonals and repeated entries) with and without GSR were 0.95 in the GSP dataset and 0.82 in the HCP dataset. However, consistent with previous work (Murphy et al., 2009; Fox et al., 2009; Weissenbacher et al., 2009), GSR reduces functional connectivity across almost all ROI pairs.

However, also consistent with previous work (Murphy et al., 2009; Gotts et al., 2013), this shift in functional connectivity was not uniform across all edges and participants. In the GSP dataset, the RSFC shift (averaged across participants) ranged from −0.58 to 0 across ROI pairs, while the RSFC shift (averaged across ROI pairs) ranged from −0.63 to −0.08 across participants. On the other hand, in the HCP dataset, the RSFC shift (averaged across participants) ranged from −0.40 to 0 across ROI pairs, while the RSFC shift (averaged across ROI pairs) ranged from −0.43 to −0.04 across participants.

#### 3.2.2 Consistent with the literature, GSR reduces motion and respiratory imaging artifacts, while strengthening distance-dependent biases

Figure 2A illustrates a widely used single-subject quality control (QC) plot pioneered by Power and colleagues (Power et al., 2014; Power, 2017). The participant was a representative mid-motion GSP subject. Median FD across all GSP participants was 0.0442mm. Mean FD of this participant was 0.0443mm. FD, DVARS and GS (before GSR) timeseries are shown in red, blue and black traces. The last two panels show the timeseries of gray matter voxels with and without GSR.

**Figure 2.**
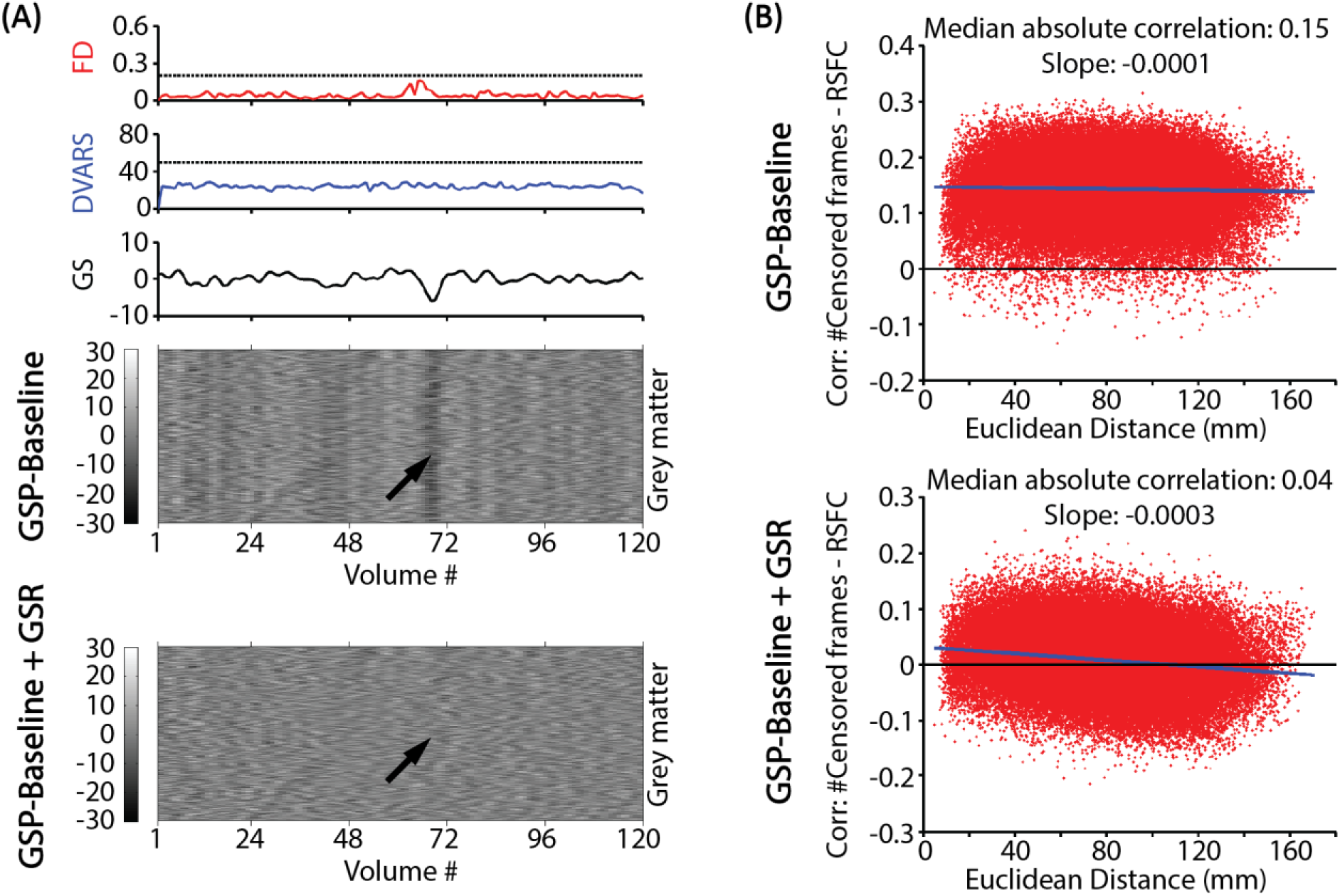
GSR reduces imaging artifacts, while exacerbating distance-dependent FC biases. (A) QC plot of a representative mid-motion GSP subject. FD (red), DVARS (blue), and GS (black) are shown in the top three panels. The horizontal lines in the first two panels indicate the thresholds used in the censoring step. The bottom two panels are signal intensity of gray matter voxels in two preprocessing pipelines: GSP-Baseline (upper) and GSP-Baseline+GSR (lower). Without GSR, motion-related global artifacts are visible in the gray matter timeseries, while GSR removes the global signal changes (black arrows). (B) Correlations between QC (number of censored frames) and RSFC are shown for two preprocessing pipelines: GSP-Baseline (upper) and GSP-Baseline+GSR (lower). Each red dot indicates one ROI pair. The x-axes are the Euclidean distances among pairs of ROIs. The functional connectivity between each pair of ROIs are correlated with the number of censored frames across all subjects and shown on the y-axes. The blue lines corresponded to linear fits of the red dots.

Consistent with previous literature (Power et al., 2014; Byrge et al., 2017; Power et al., 2018), without GSR, there were multiple bands of signal increase or decrease across most brain voxels that extended for many time points. A particularly salient global decrease in signal intensity appeared right after a peak in FD, which was mostly attenuated by GSR (black arrow in Figure 2A). Similar observations could be made for a mid-motion HCP participant (Figure S2A).

Figure 2B shows another common type of QC-FC plot with and without GSR for the GSP dataset. Each dot represents an ROI pair. The x-axis corresponds to the distance between the pair of ROIs. The y-axis corresponds to the correlation between the FC of the ROI pair and QC (number of censored frames) across participants. Consistent with previous work (Power et al., 2014; Burgess et al., 2016; Ciric et al., 2017), the median absolute FC-QC correlations decreased from 0.15 to 0.04 after GSR, suggesting an improvement in data quality with GSR. Similar results were obtained with the HCP dataset (Figure S2B).

Furthermore, replicating previous work, GSR exacerbated the distance-dependent biases in QC-FC correlations. In the case of the GSP dataset (Figure 2B), GSR introduces a negative bias to distance-dependent QC-FC correlations with slope decreasing from −0.0001 to −0.0003, consistent with previous work (Power et al., 2014; Burgess et al., 2016; Ciric et al., 2017). However, in the HCP dataset (Figure S2B), GSR introduces a positive bias to distance-dependent QC-FC correlations with slope increasing from 0.0001 to 0.0002.

### 3.3 GSR increases behavioral variance explained by RSFC

#### 3.3.1 Behavioral-RSFC association in the GSP dataset

Figure 3A shows the behavioral variance explained by RSFC averaged across all 23 behavioral measures in the GSP dataset. The blue and green bars represent the interquartile range (IQR) across jackknife samples. The non-overlapping IQRs suggest a robust increase in explained behavioral variance with GSR. The mean improvement (averaged across all behavioral measures and jackknife samples) was 46.6%.

**Figure 3.**
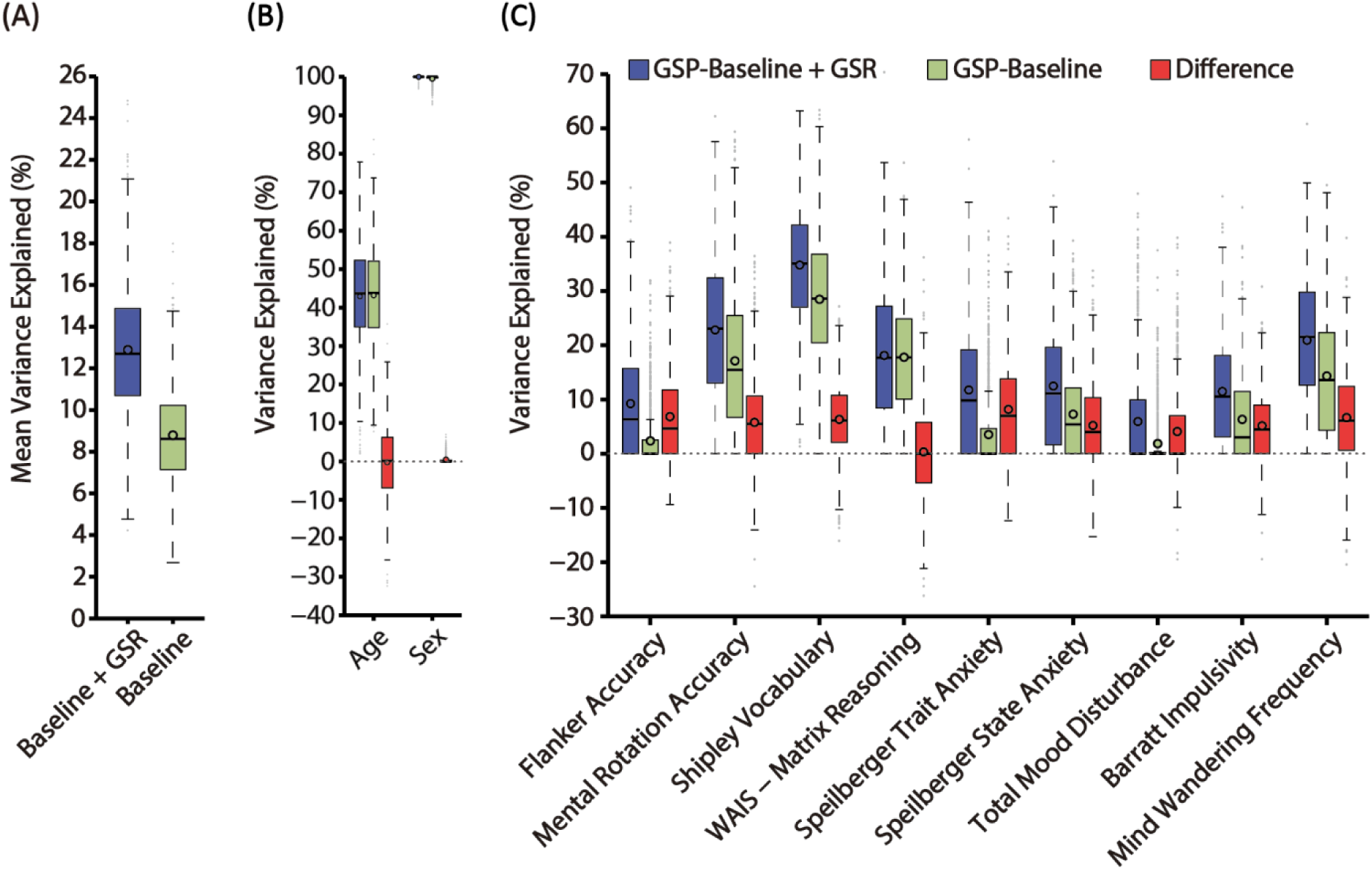
GSR improves behavioral variance explained by RSFC in the GSP dataset using the variance component model. (A) Behavioral variance explained by RSFC averaged across all 23 behaviors for two preprocessing pipelines: GSP-Baseline+GSR (blue) and GSP-Baseline (green). The “boxes” show the median and interquartile range (IQR) of explained variance across all jackknife samples. The whisker length is 1.5 IQR. Black circles indicate mean. Outliers are shown by grey dots. (B) Variance explained by RSFC for age and sex. (C) Behavioral variance explained by RSFC for 9 behavioral measures. For each behavioral measure, the explained variance by the GSP-Baseline+GSR pipeline, GSP-Baseline pipeline, and difference between the two pipelines are shown in blue, green and red respectively.

Figure 3B shows the RSFC-explained variance for age and sex. Figure 3C shows the RSFC-explained variance for 9 behavioral measures in the GSP dataset. Figure S3 shows the RSFC-explained variance for the remaining 14 behavioral measures. Like Figure 3A, the blue and green boxplots correspond to the GSP-Baseline+GSR and GSP-Baseline preprocessing pipelines respectively. Differences between the two pipelines are illustrated by the red boxplots. For 15 of the 25 demographic and behavioral measures (Figures 3B, 3C and S3), the red boxes were entirely above zero, implying that for the 15 measures, GSR increases the explained variance in at least 75% of the jackknife samples. For 22 measures, GSR increased the explained behavioral variance, although the red “boxes” were not entirely above zero (e.g., Reward Dependence). Across the 9 behavioral measures (Figure 3C), only WAIS – Matrix Reasoning did not show a robust increase in RSFC-explained variance after GSR. Across all 25 measures, there was only one behavioral measure (Behavioral Activation - Reward) for which the baseline processing pipeline (non-GSR) exhibited greater explained variance in at least 75% of the jackknife samples. Thus, the overall increase in behavior-RSFC association (Figure 3A) was not driven by specific behavioral measures, but was the result of robust improvements across most of the behavioral measures examined.

#### 3.3.2 Behavioral-RSFC association in the HCP dataset

Figure 4A shows behavioral variance explained by RSFC averaged across 58 behavioral measures in the HCP dataset. Similar to before, the blue and green boxplots corresponded to HCP-Baseline+GSR and HCP-Baseline respectively. The non-overlapping IQRs suggest a robust increase in explained behavioral variance with GSR. Compared to baseline, the mean explained behavioral variance increased by 40.4% with GSR.

**Figure 4.**
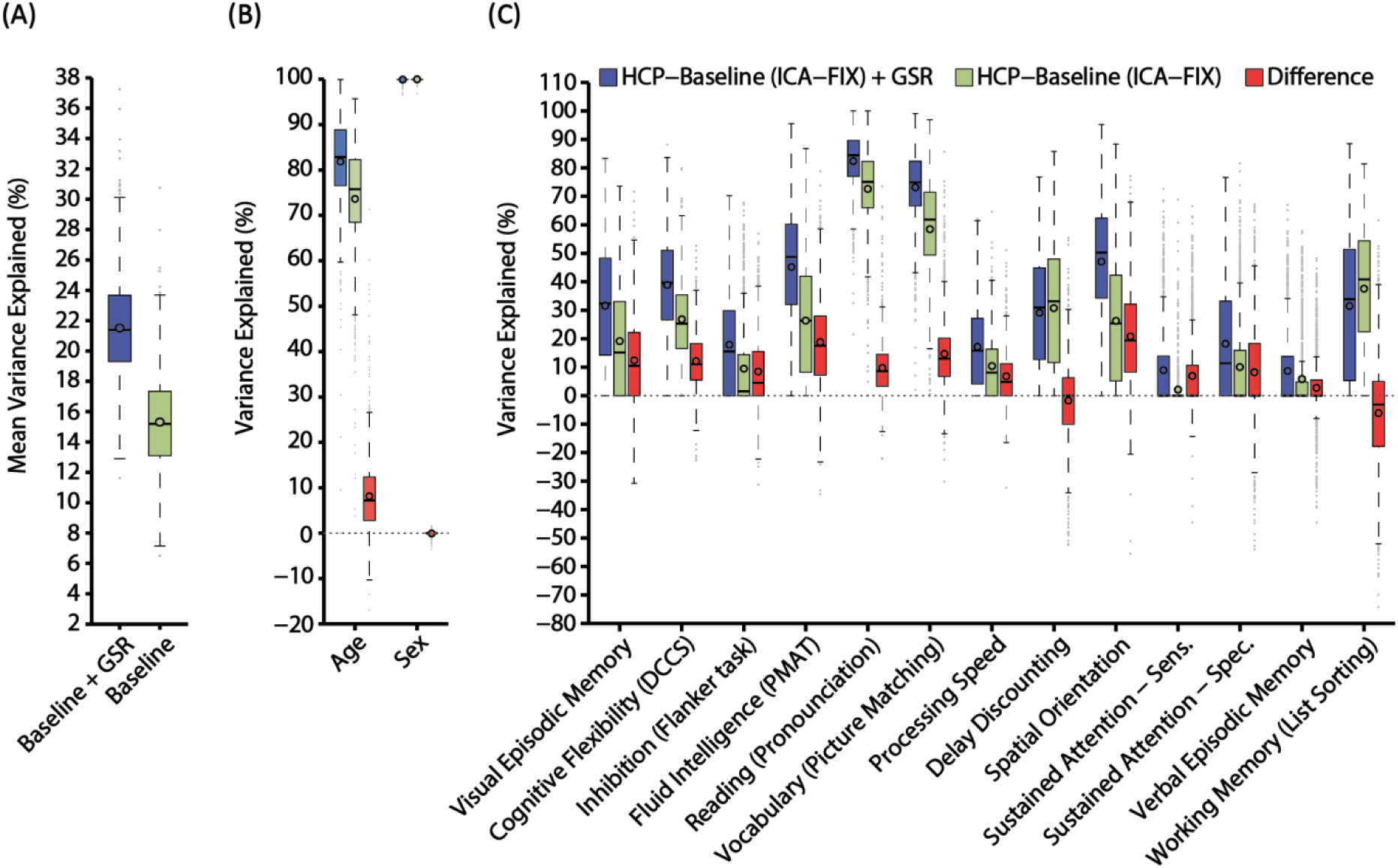
GSR improves behavioral variance explained by RSFC in the HCP dataset using the variance component model. (A) Behavioral variance explained by RSFC averaged across all 58 behaviors for two preprocessing pipelines: HCP-Baseline+GSR (blue) and HCP-Baseline (green). The “boxes” show the median and interquartile range (IQR) of explained variance across all jackknife samples. The whisker length is 1.5 IQR. Black circles indicate mean. Outliers are shown by grey dots. (B) Variance explained by RSFC for age and sex. (C) Behavioral variance explained by RSFC for 13 cognitive measures. For each behavioral measure, the explained variance by the HCP-Baseline+GSR pipeline, HCP-Baseline pipeline, and difference between the two pipelines are shown in blue, green and red respectively.

Figure 4B shows the RSFC-explained variance for age and sex. Figures 4C, S4, and S5 show the RSFC-explained behavioral variance for individual behavioral measures. Figure 4C contains 13 HCP cognitive measures. Figures S4 and S5 contain the remaining 45 behavioral measures, including alertness, motor, sensory, personality, emotion, and other in-scanner measures. Like before, the blue and green boxplots correspond to the HCP-Baseline+GSR and HCP-Baseline preprocessing pipelines respectively. Differences between the two pipelines are illustrated by the red boxplots.

For 40 of the 60 measures, the red “boxes” were entirely above zero, implying that for the 40 measures, GSR increases the explained variance in at least 75% of the jackknife samples. For 47 measures, GSR increased the explained behavioral variance, although the red “boxes” were not entirely above zero (e.g., Walking Memory (N-back)). Conversely, there were 5 behavioral measures for which the baseline processing pipeline (non-GSR) exhibited greater explained variance in at least 75% of the jackknife samples.

For the 13 cognitive measures in Figure 4C, except for Delay Discounting and Working Memory (List Sorting), all cognitive measures have higher RSFC-explained variance after GSR. Of the 23 social-emotional measures (Figure S5), the behavioral variance explained by RSFC decreased after GSR for 5 measures (Emot. Recog. – Happy, Emot. Recog. – Sad, Anger – Hostility, Fear – Somatic Arousal, and Self - Efficacy). Thus, the overall increase in behavior-RSFC association (Figure 4A) was not driven by specific behavioral measures, but was the result of robust improvements across most of the behavioral measures examined.

It is worth noting that the behavioral variance explained by RSFC was higher in the HCP dataset compared with the GSP dataset. One reason could be the significantly greater amount of data per participant in the HCP dataset (1 hour) compared with the GSP dataset (at most 12 minutes).

### 3.4 GSR improves RSFC-based behavioral prediction accuracies

#### 3.4.1 RSFC prediction of behavior in the GSP dataset

Figure 5 shows the RSFC-based prediction accuracies with and without GSR in the GSP dataset. Figure 5A shows the mean prediction accuracy averaged across 23 behavioral measures. Like previous figures, the blue and green boxplots correspond to GSP-Baseline+GSR and GSP-Baseline preprocessing pipelines respectively with the bars representing the IQR across random cross-validation splits. The non-overlapping IQRs suggest a robust increase in behavioral prediction accuracies with GSR. The mean improvement was 64.3%.

**Figure 5.**
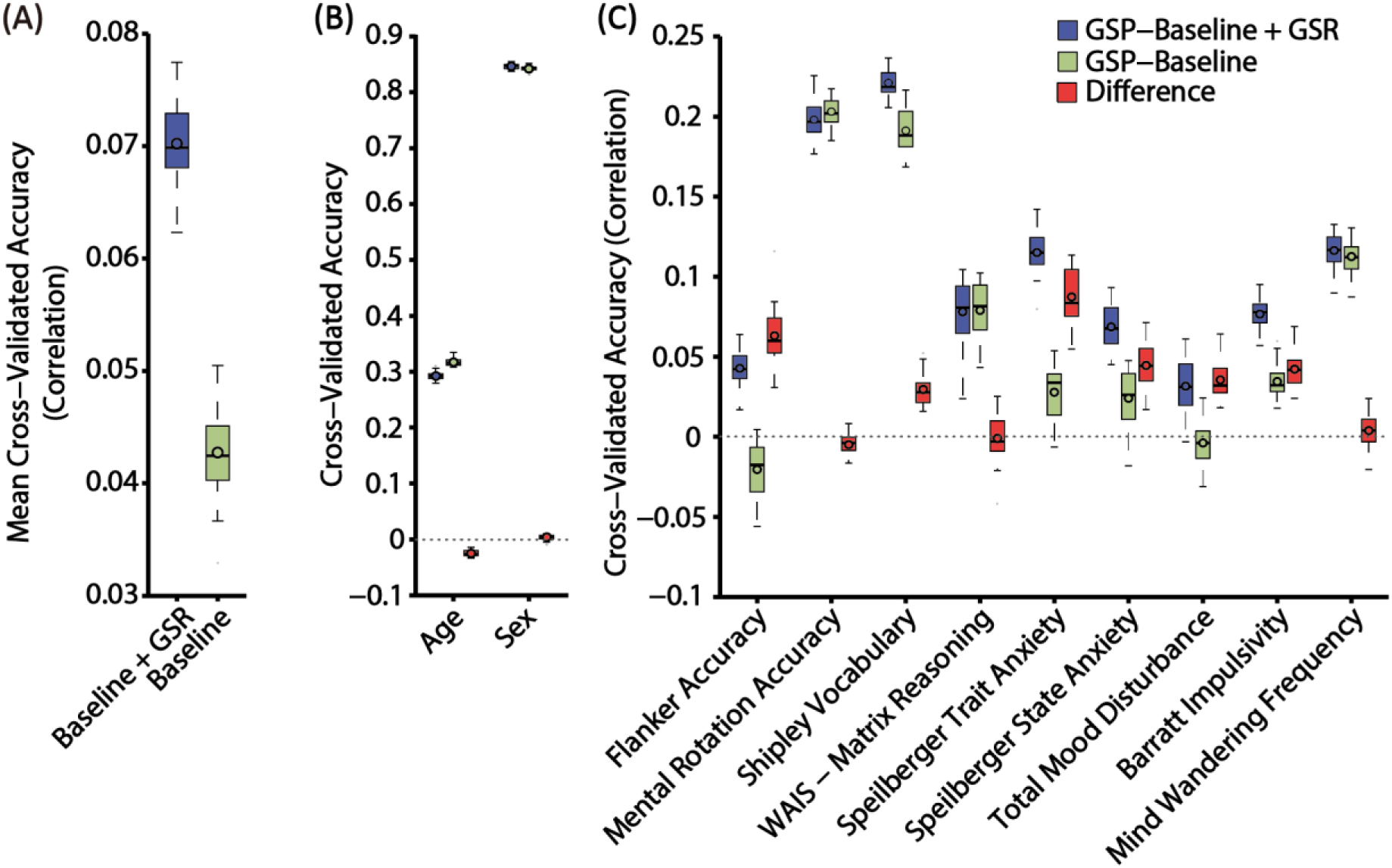
GSR improves RSFC-based behavioral prediction accuracies in the GSP dataset using kernel ridge regression. (A) Test accuracies averaged across all 23 behaviors for two preprocessing pipelines: GSP-Baseline+GSR (blue) and GSP-Baseline (green). The “boxes” show the median and interquartile range (IQR) of test accuracies across 20 random cross-validation splits. The whisker length is 1.5 IQR. Black circles indicate mean. Outliers are shown by grey dots. (B) Test accuracies for age and sex. (C) Test accuracies for 9 behavioral measures. For all measures, the accuracies of the GSP-Baseline+GSR pipeline, GSP-Baseline pipeline, and difference between the two pipelines are shown in blue, green and red respectively.

Figures 5B shows the prediction accuracies for age and sex. Figure 5C shows the prediction accuracies for 9 behavioral measures in the GSP dataset. Figure S6 shows the prediction accuracies for the remaining 14 behavioral measures. Like before, the blue and green boxplots correspond to the GSP-Baseline+GSR and GSP-Baseline preprocessing pipelines respectively. Differences between the two pipelines are illustrated by the red boxplots.

The red boxes were entirely above zeros for 15 of the 25 demographic and behavioral measures (Figures 5B, 5C, and S6), implying that for the 15 measures, GSR increased the prediction accuracies in at least 75% of the random cross-validation splits. Conversely, for 4 of the total 25 measures, baseline preprocessing pipeline (non-GSR) exhibited greater prediction accuracy in at least 75% of the random data splits. Thus, the overall increase in behavioral prediction accuracy (Figure 5A) was not driven by specific measures, but was the result of robust improvements across many of the demographic and behavioral measures examined.

#### 3.4.2 RSFC prediction of behavior in the HCP dataset

Figure 6 shows the RSFC-based prediction accuracies with and without GSR in the HCP dataset. Figure 6A shows the mean prediction accuracy averaged across 58 behavioral measures. Like previous figures, the blue and green boxplots correspond to HCP-Baseline+GSR and HCP-Baseline preprocessing pipelines respectively. The non-overlapping IQRs suggest a robust increase in behavioral prediction accuracies with GSR. The mean improvement was 11.6% using kernel regression.

**Figure 6.**
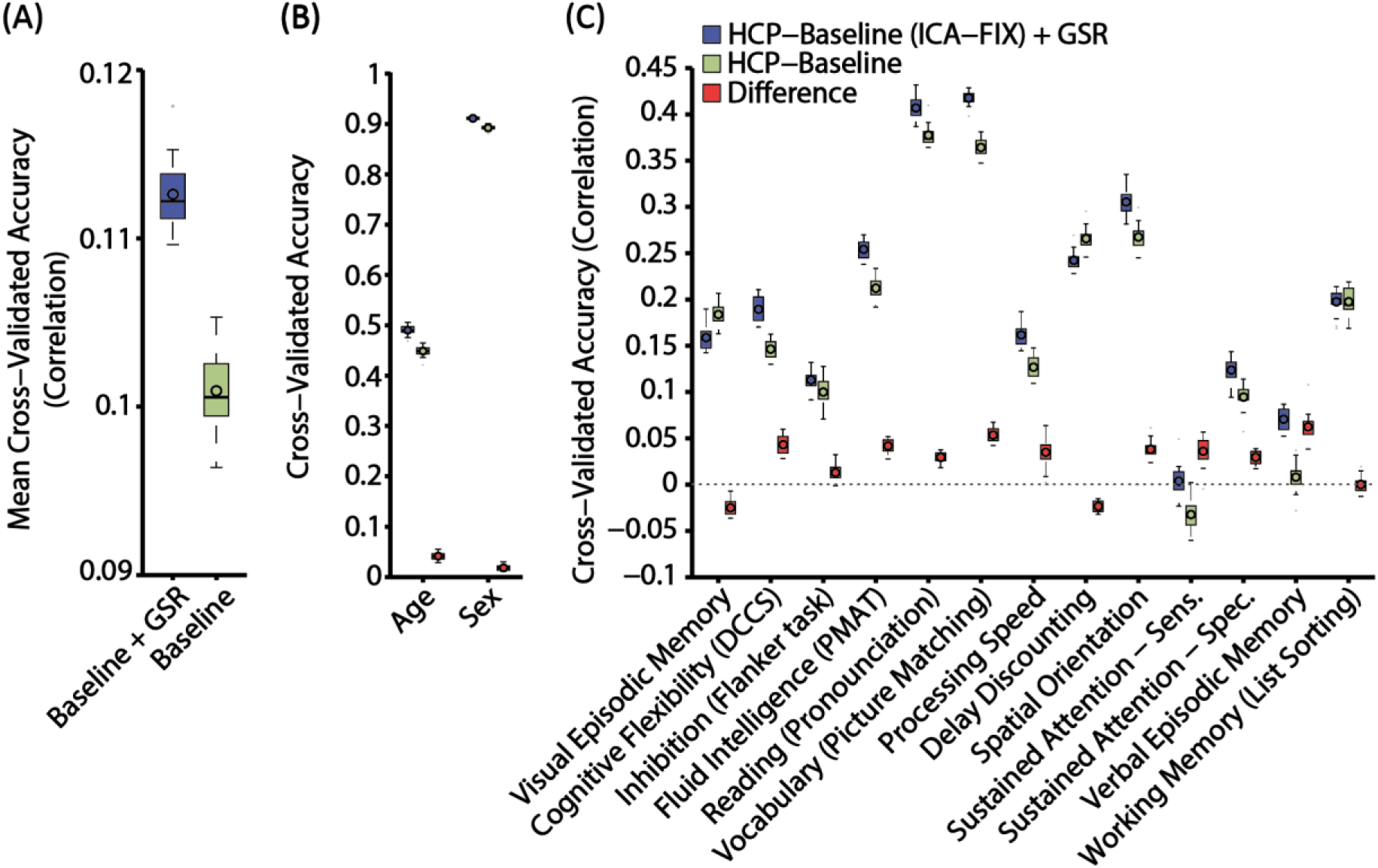
GSR improves RSFC-based behavioral prediction accuracies in the HCP dataset using kernel ridge regression. (A). Test accuracies averaged across all 58 behaviors for two preprocessing pipelines: HCP-Baseline+GSR (blue) and HCP-Baseline (green). The “boxes” show the median and interquartile range (IQR) of test accuracies across 20 random cross-validation splits. The whisker length is 1.5 IQR. Black circles indicate mean. Outliers are shown by grey dots. (B). Test accuracies for age and sex. (C). Test accuracies for 13 cognitive measures. For each behavioral measure, the accuracies of the HCP-Baseline+GSR pipeline, HCP-Baseline pipeline, and difference between the two pipelines are shown in blue, green and red respectively.

Figures 6B shows the prediction accuracies for age and sex. Figure 6C shows the prediction accuracies for 13 HCP cognitive measures. Figures S7 and S8 show the prediction accuracies for the remaining 45 behavioral measures. Like before, the blue and green boxplots correspond to the HCP-Baseline+GSR and HCP-Baseline preprocessing pipelines respectively. Differences between the two pipelines are illustrated by the red boxplots.

The red boxes were entirely above zero for 33 of the 60 demographic and behavioral measures (Figures 6B, 6C, S7, and S8), implying that for the 33 measures, GSR increased the prediction accuracy in at least 75% of the random cross-validation splits. Conversely, there were 14 behavioral measures for which the baseline preprocessing pipeline (non-GSR) exhibited greater prediction accuracies in at least 75% of the random data splits. Thus, the overall increase in behavioral prediction accuracy (Figure 6A) was not driven by specific measures, but was the result of robust improvements across many of the behavioral measures examined.

### 3.5 Consistencies and heterogeneities across approaches and behavioral measures

#### 3.5.1 Consistencies between RSFC-behavioral associations and RSFC-behavioral prediction accuracies

To investigate whether RSFC-behavioral associations and RSFC-behavioral prediction accuracies are consistent, Figure 7 shows scatterplots of RSFC-explained behavioral variance (variance component model) and prediction accuracies (kernel ridge regression). Each dot in each plot represents a single behavioral measure with corresponding prediction accuracy and RSFC-explained variance. For both datasets and both pipelines, the Pearson’s correlations between the RSFC-explained behavioral variance and behavioral prediction accuracies were high, ranging from 0.75 to 0.87. In other words, if a behavioral measure was well explained by RSFC using the variance component model, it was also better predicted using kernel ridge regression.

**Figure 7.**
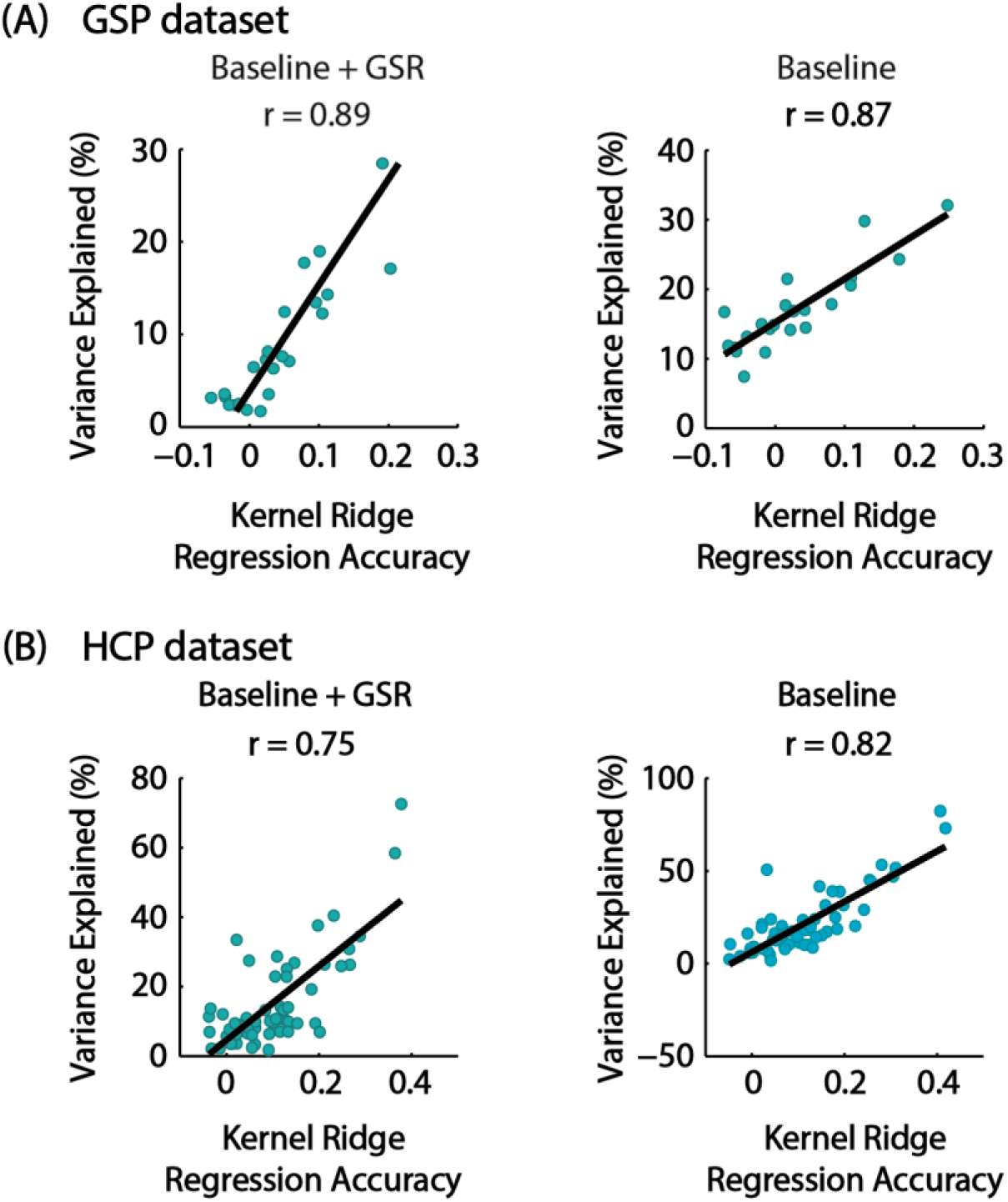
Consistency across explained behavioral variance and kernel ridge regression accuracies. (A) Scatterplots of explained behavioral variance (variance component model) and kernel ridge regression accuracies for 23 GSP behavioral measures with and without GSR. (B) Scatterplots of explained behavioral variance (variance component model) and kernel ridge regression accuracies for 58 HCP behavioral measures with and without GSR. Each blue dot represents a behavioral measure. Black lines represent the linear fit of blue dots. Pearson’s correlation coefficients between explained variance and accuracies are reported.

Furthermore, GSR-related improvements in RSFC-explained behavioral variance and RSFC-based behavioral prediction accuracies were modestly correlated across behavioral measures in both the GSP (r = 0.40) and HCP (r = 0.49) datasets (Figure 8). In other words, a behavioral measure with a larger improvement in RSFC-explained variance after GSR will also enjoy greater improvement in prediction accuracy after GSR.

**Figure 8.**
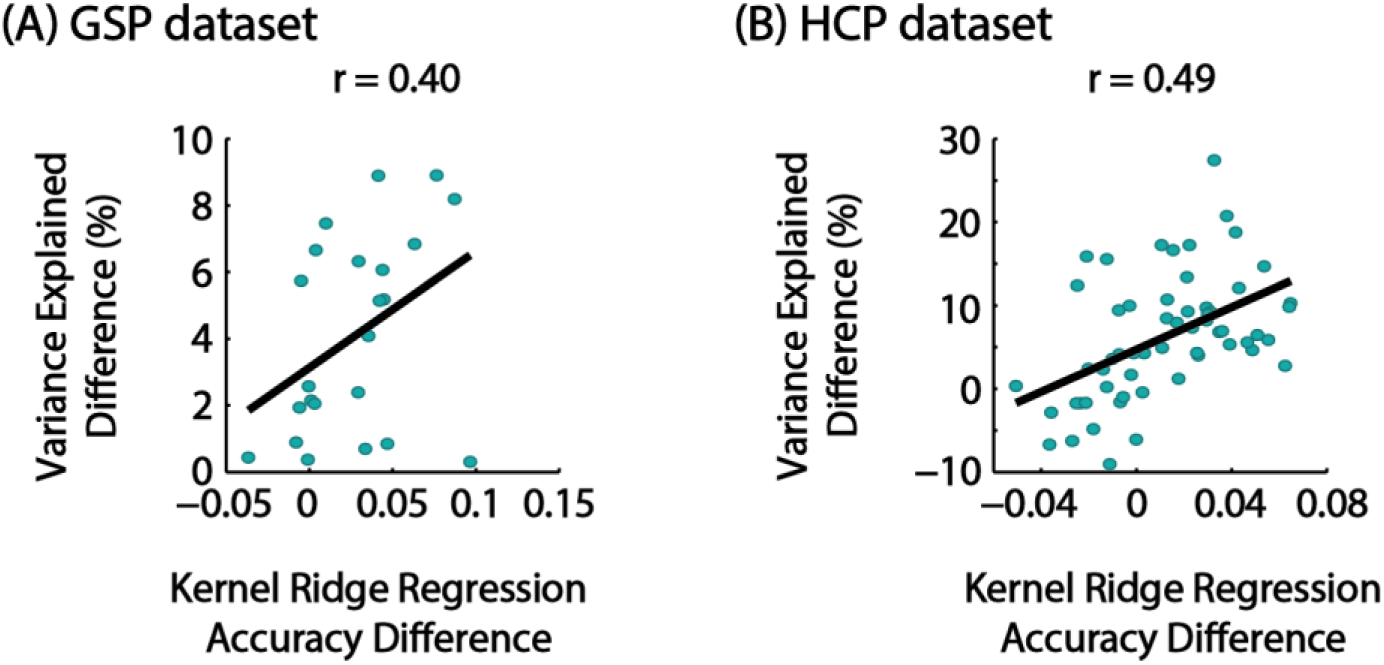
Consistencies in GSR-related improvements between RSFC-explained behavioral variance and kernel ridge regression accuracies. (A) Scatterplots of GSR-related change in RSFC-explained behavioral variance and GSR-related change in kernel ridge regression accuracies for 23 GSP behavioral measures. (B) Scatterplots of GSR-related change in RSFC-explained behavioral variance and GSR-related change in kernel ridge regression accuracies for 58 HCP behavioral measures. Each blue dot represents a behavioral measure. Black lines represent the linear fit of blue dots. Pearson’s correlation coefficients between explained variance and accuracies are reported.

#### 3.5.2 Relationship with baseline associations and prediction accuracies

In the case of the variance component model, GSR-related improvements in RSFC-explained variance were weakly correlated with the RSFC-explained variance of the baseline preprocessing pipelines for both the GSP (r = 0.23) and HCP (r = 0.24) datasets.

In the HCP dataset, GSR-related improvements in behavioral prediction accuracies were also weakly correlated with the prediction accuracies of the baseline preprocessing pipeline (r = 0.10). On the other hand, in the GSP dataset, GSR-related improvements in behavioral prediction accuracies were moderately negatively correlated with the prediction accuracies of the baseline preprocessing pipeline (r = −0.51). In other words, if a behavioral measurement was poorly predicted with the baseline preprocessing pipeline, then the GSR-related improvement in behavioral prediction would be greater.

#### 3.5.3 GSR preferentially improves RSFC-explained behavioral variance and behavioral prediction accuracies for task performance measures

Figure 9A shows the mean GSR-related improvements in RSFC-explained variance averaged across 27 HCP task-performance measures (green) and 24 self-reported measures (red). On average, task-performance measures enjoyed greater GSR-related improvement (7.9%) compared with self-reported measures (4.4%). Figure 9B shows the GSR-related improvements in RSFC-explained variance for individual behavioral measures sorted by the magnitude of improvement.

**Figure 9.**
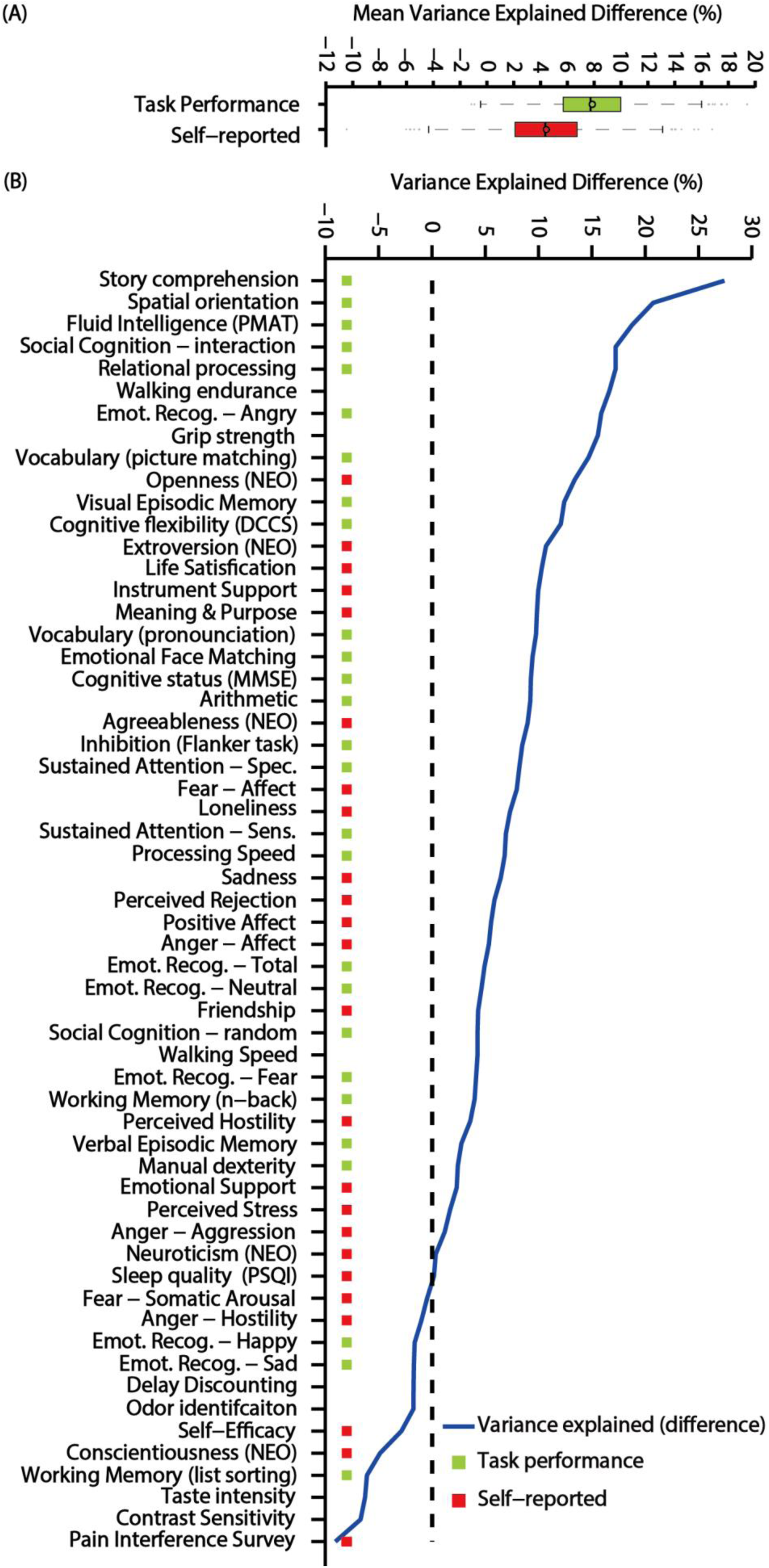
GSR preferentially improves RSFC-explained behavioral variance for task-performance measures compared with self-reported measures. (A) Improvement in RSFC-explained behavioral variance averaged across all HCP task-performance measures (green) and across all self-reported measures (red). The “boxes” show the median and interquartile range (IQR) of explained variance across all jackknife samples. The whisker length is 1.5 IQR. Black circles indicate mean. (B) Behavioral measures are ordered based on the improvement in RSFC-explained behavioral variance (blue line). Behavioral measures marked with green color are considered task-performance measures. Behavioral measures marked with red color are considered self-reported measures.

Figure S9A shows the mean improvement in GSR-related behavioral prediction accuracies averaged across task performance measures (green) and self-reported measures (red). On average, task-performance measures enjoyed greater GSR-related improvement (0.016) compared with self-reported measures (0.0084). Figure S9B shows the GSR-related improvements in behavioral prediction accuracies for individual behavioral measures sorted by the magnitude of improvement.

### 3.6 Further motion analyses

#### 3.6.1 GSR improves RSFC-explained variance even without including FD & DVARS as covariates in the variance component model

Figure 10A shows the behavioral variance explained by RSFC averaged across 23 behavioral measures in the GSP dataset for both processing pipelines, with and without including FD and DVARS as nuisance covariates in the variance component model. Interestingly, the variance explained were unchanged regardless of whether including FD and DVARS were included as covariates.

**Figure 10.**
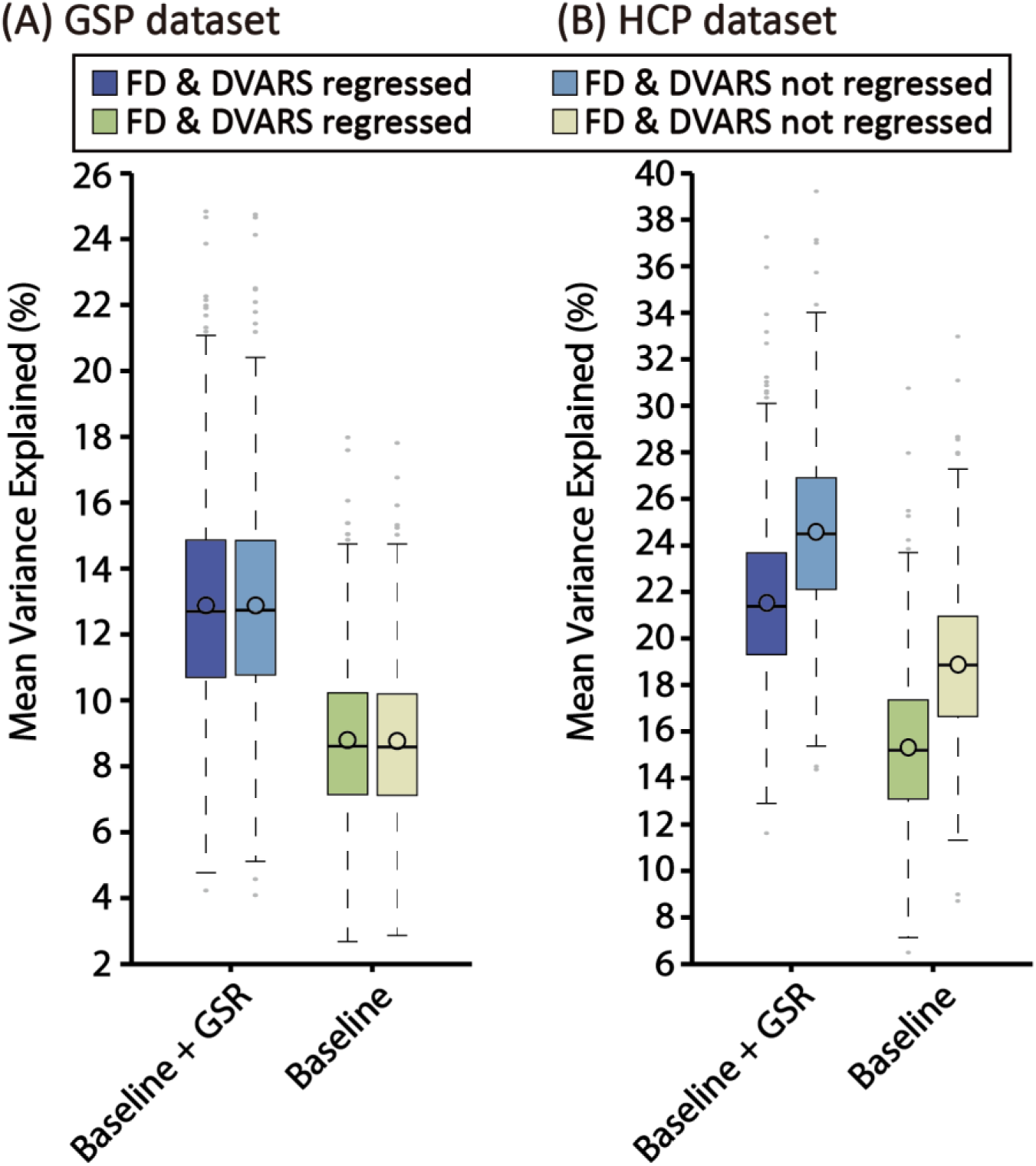
GSR improves RSFC-explained variance in both GSP and HCP datasets even when FD and DVARS were not included as nuisance covariates in the variance component model. (A) GSP dataset. (B) HCP dataset. Blue boxplots represent the variance explained by RSFC averaged across all behaviors with the Baseline+GSR preprocessing pipeline. Green boxplots represent the variance explained by RSFC averaged across all behaviors with the Baseline preprocessing pipeline. Darker colors represent results when FD and DVARS were included as covariates; lighter colors represent the results when FD and DVARS were not regressed. The “boxes” show the median and interquartile range (IQR) of explained variance across all jackknife samples. The whisker length is 1.5 IQR. Black circles indicate mean. Outliers are shown by grey dots.

Figure 10B shows the behavioral variance explained by RSFC averaged across 58 behavioral measures in the HCP dataset for both processing pipelines, with and without including FD and DVARS as nuisance covariates in the variance component model. For both pipelines, the explained variance was higher when FD and DVARS were not included as nuisance covariates. Nevertheless, the explained variance was still higher for the baseline+GSR pipeline than the baseline pipeline.

#### 3.6.2 RSFC-explained behavioral variance cannot be explained by motion

Tables S3 and S4 show the correlations of behavioral measures with FD (and DVARS) in the GSP and HCP datasets respectively. Figure 11A shows that the mean behavioral variances explained by RSFC in the GSP dataset for both processing pipelines were much greater than the absolute correlation between the behavioral measures and FD (expressed in terms of percentage variance), as well as the absolute correlation between the behavioral measures and DVARS (expressed in terms of percentage variance). The same was true for the HCP dataset (Figure 11B). Therefore, RSFC-explained behavioral variance for both baseline and baseline+GSR processing pipelines cannot be simply due to motion.

**Figure 11.**
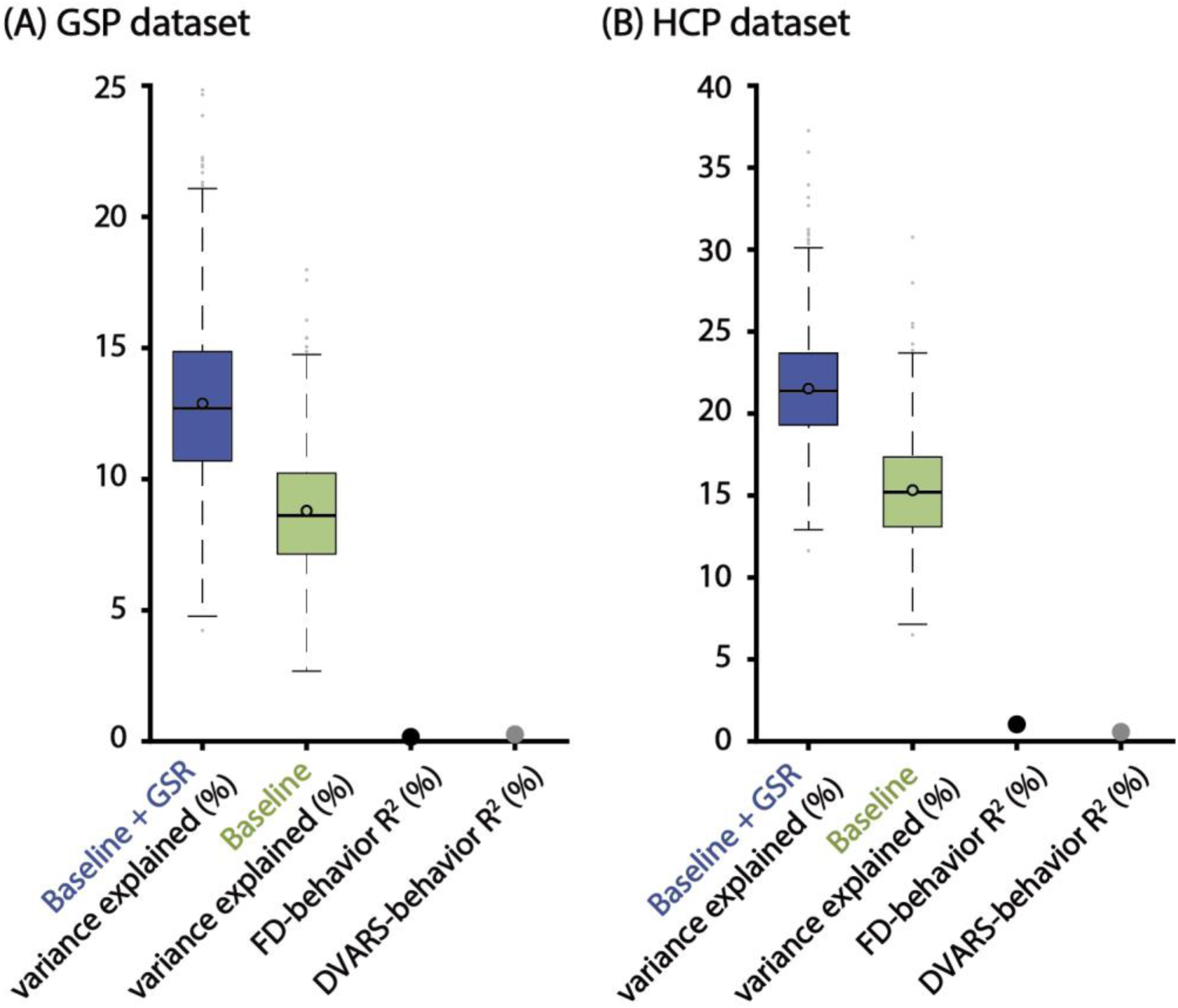
RSFC-explained behavioral variance cannot be simply due to motion in the (A) GSP and (B) HCP datasets. Boxplots: variance explained by RSFC averaged across all behavioral measures. Black dots: correlation between motion (FD or DVARS) and behavior (expressed in terms of percentage variance) averaged across all behavioral measures. The “boxes” show the median and interquartile range (IQR) across 1000 jackknife samples. The whisker length is 1.5 IQR. Black circles indicate mean.

#### 3.6.3 RSFC-based behavioral prediction accuracies cannot be explained by motion

Figure 12A shows that the mean RSFC-based behavioral prediction accuracy in the GSP dataset for the baseline+GSR processing pipeline was greater than the mean absolute correlation between the behavioral measures and FD, as well as the mean absolute correlation between the behavioral measures and DVARS. However, in the case of the baseline processing pipeline, the absolute correlation between the behavioral measures and DVARS was bigger than the prediction accuracy.

**Figure 12.**
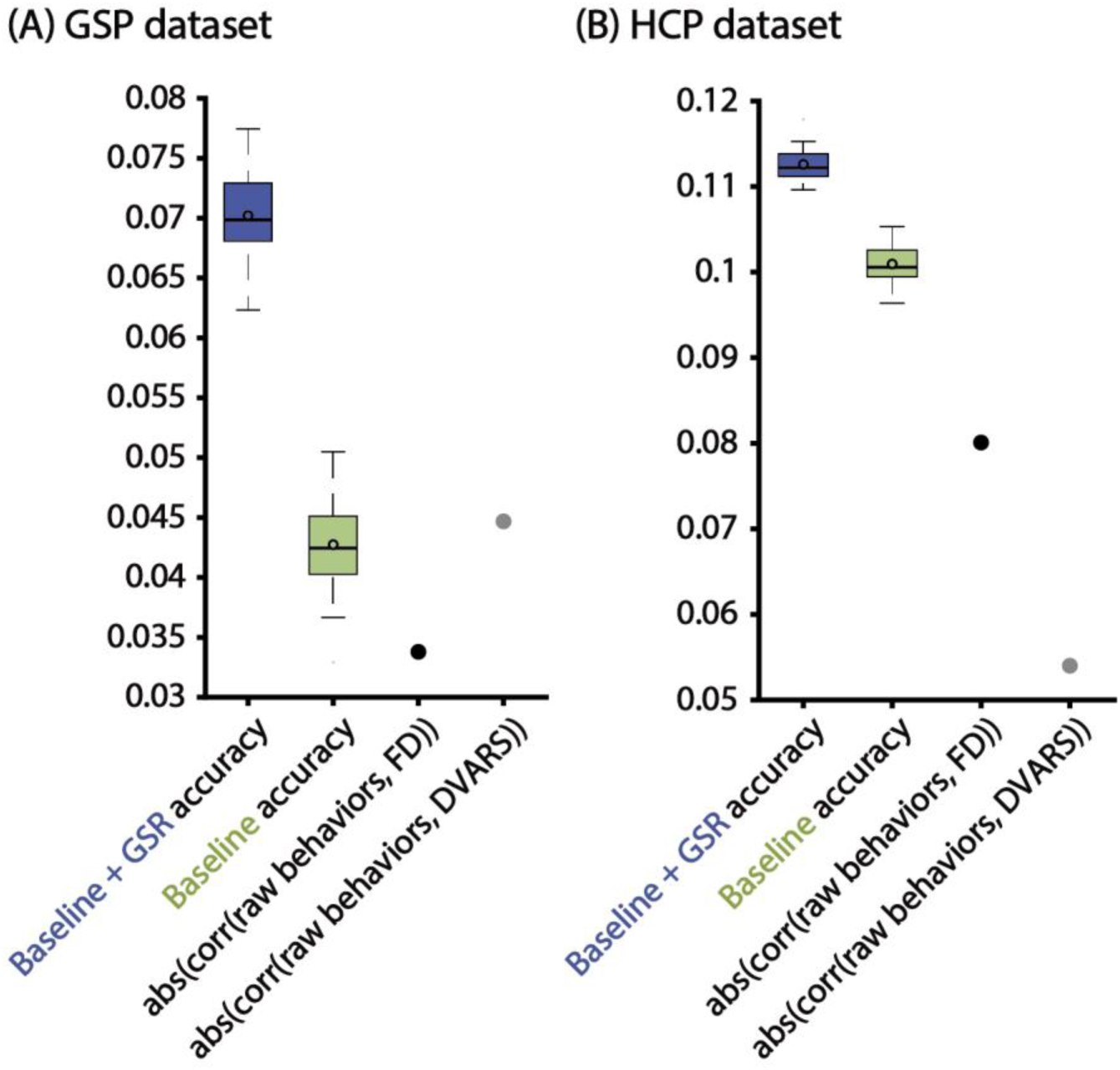
RSFC-based behavioral prediction accuracies cannot be simply due to motion in the (A) GSP and (B) HCP datasets. Boxplots: RSFC-based prediction accuracies averaged across all behavioral measures. Black dots: absolute correlation between motion (FD or DVARS) and behavior averaged across all behavioral measures. The “boxes” show the median and interquartile range (IQR) across 1000 jackknife samples. The whisker length is 1.5 IQR. Black circles indicate mean.

On the other hand, Figure 12B shows that the average RSFC-based behavioral prediction accuracies in the HCP dataset for both processing pipelines were greater than the absolute correlation between the measures and FD, as well as the absolute correlation between the measures and DVARS.

Overall, RSFC-based behavioral prediction accuracies for the baseline+GSR preprocessing pipeline cannot be simply due to motion in both GSP and HCP datasets. In the case of the baseline preprocessing pipeline, this was only true for the HCP dataset.

## 4. Discussion

Using the variance component model, we showed that GSR strengthened the association between behavior and RSFC in young healthy adults from two large-scale datasets. Of the 81 behavioral measures examined, there were robust (>75% jackknife samples) improvements for 53 measures, with only 6 measures in the opposite direction. The improvements were substantial: average increase of 47% in the GSP dataset and average increase of 40% in the HCP dataset. Using kernel ridge regression, we found that GSR improved RSFC-based behavioral prediction accuracies. 45 of the 81 behavioral measures showed robust (>75% random cross-validation splits) improvements with 17 measures in the opposite direction. The improvements were substantial in the GSP dataset, but modest in the HCP dataset: average improvement of 64% in the GSP dataset and 12% improvement in the HCP dataset.

Therefore, across both methods and datasets, we found that GSR improved behavioral associations and prediction accuracies. Furthermore, both approaches were consistent in the sense that a behavioral measure well explained by RSFC using the variance component model was also better predicted by kernel ridge regression (Figure 7). In addition, a behavioral measure with a larger improvement in RSFC-explained variance after GSR also tended to enjoy greater improvement in prediction accuracy after GSR (Figure 8). Further analysis suggests that task performance measures appeared to benefit more from GSR than self-reported measures (Figures 9 and S9).

### Interpretations and implications

We note that our results could have been in the opposite direction. After all, it is well known that the global signal contains neural information (Schölvinck et al., 2010; Matsui et al., 2016), so regressing the global signal might potentially reduce the associations between RSFC and behavioral traits. On the other hand, GSR is known to reduce imaging artifacts (Satterthwaite et al., 2013; Power et al., 2014; Burgess et al., 2016; Ciric et al., 2016; Parkes et al., 2018), so using the cleaner data might strengthen RSFC-behavior association. Yet another consideration is that because certain behavioral traits (e.g., reading pronunciation) are correlated with motion (Siegel et al., 2017), more effective removal of motion-related artifacts via GSR might weaken the RSFC-behavior association (which might be desirable).

Given that our experiments suggest that GSR strengthens RSFC-behavior association (on average), one possibility is that the neural component within the global signal might not be associated with the examined behavioral traits. For example, the neural component might reflect transient neural activity, which might not relate to stable behavioral traits measured outside the scanner. A second possibility is that having cleaner data (from removing the noise component contained in the global signal) outweighs any negative effect from losing the neural component embedded in the global signal, so that overall, GSR improves behavior-RSFC association. Dissociating these two possibilities might not be easy, but a recent study suggested that GSR reduced task-relevant information in task-fMRI, and instead advocated the use of temporal ICA to remove global artifacts, while retaining neural information (Glasser et al., 2018). Thus, we hope to evaluate the effects of temporal ICA on behavior-RSFC association in the future.

Given that GSR is more effective at removing global motion and respiratory artifacts than baseline processing (Figures 2; S2; Burgess et al., 2016; Ciric et al., 2017), the improved explained behavioral variance and prediction accuracies (with GSR) were not confounded by imaging-related artifacts. If the reverse were true (i.e., non-GSR explained greater behavioral variance and improved behavioral prediction accuracies compared with GSR), the interpretation would be considerably trickier because it would be hard to rule out the possibility that the greater explained behavioral variance might be due to imaging artifacts correlated with behavioral measurements (Siegel et al., 2017).

The variance component model and the kernel ridge regression were set up, so that inter-subject similarity was defined based on the correlation of whole-brain RSFC matrices. This choice was motivated by seminal work on functional connectivity fingerprints (Finn et al., 2015). Work from our lab also suggests that this particular choice of inter-subject similarity yields competitive behavioral prediction accuracy (He et al., 2018). We note that our experiments with linear ridge regression (not shown) yielded similar conclusions to the variance component model and kernel ridge regression, suggesting that the benefits of GSR were not limited to this particular choice of similarity metric.

### Limitations, methodological considerations and future work

Because our study is focused on young healthy adults, our conclusions might not hold for other cohorts. Indeed, some studies have suggested that GSR weakens differences between healthy controls and diseased populations, such as autism spectrum disorder (Gotts et al., 2013) and schizophrenia (Yang et al., 2014). On the other hand, Parkes and colleagues showed that group differences between healthy controls and schizophrenia were only seen with GSR (Parkes et al., 2018). However, given that GSR is known to more effectively remove global motion and respiratory artifacts compared with other de-noising approaches (Ciric et al., 2017; Power et al., 2017b), it is not easy to rule out that any group differences without GSR were not due to imaging artifacts. Nevertheless, given the availability of large-scale datasets of diseased and lifespan cohorts (Milham et al., 2012; Di Martino et al., 2014; Poldrack et al., 2016), we hope to extend the current study to lifespan and diseased datasets.

Previous work on rs-fMRI behavioral prediction has suggested that whole-brain RSFC would more effectively explain behavioral variance than a more restricted set of networks (Finn et al., 2015; Kong et al., 2018), so our analyses were limited to whole brain RSFC. However, a drawback of defining inter-subject similarity based on the whole-brain RSFC matrices was that we were unable to distinguish whether certain connections (e.g., within specific networks or between certain networks) were more important for explaining behavioral variance. Furthermore, it remains unclear whether GSR improves RSFC-behavioral associations for specific connections or networks. Therefore, future work might involve extending this study to investigate the effect of GSR on RSFC-behavior association for specific networks (Nostro et al., 2018) or at the voxel level (Shehzad et al., 2014; Gong et al, 2018).

Despite the effectiveness of GSR in removing imaging artifacts (Ciric et al., 2017), there were likely residual artifacts. For example, when FD and DVARS were not included as nuisance covariates in the HCP dataset, the explained behavioral variances increased by about the same amount for both baseline and baseline+GSP pipelines (Figure 10B), suggesting the presence of residual artifacts for both pipelines. Interestingly, excluding FD and DVARS as nuisance covariates has almost no impact on explained behavioral variances in the GSP dataset (Figure 10A). One possible explanation is that the behavioral measures were more correlated with motion in the HCP dataset. Indeed, if we only considered 29 of the 58 HCP behavioral measures that were the least associated with FD, then including or excluding FD and DVARS as nuisance covariates has almost no impact on explained behavioral variances in the HCP dataset. Nevertheless, the good news is that in both GSP and HCP datasets, both RSFC-explained behavioral variance and RSFC-based prediction accuracies after GSR were much greater than the magnitude of correlations between motion (FD/DVARS) and behavior (Figures 11 and 12), suggesting that motion cannot fully account for the explained behavioral variance and prediction accuracies.

Finally, it is worth noting that while the preprocessing pipelines considered in this study are representative of most published papers, the exact processing choices are variable across the literature, so we have taken the pragmatic approach of being consistent with our previous work (Yeo et al., 2011; Yeo et al., 2015a; He et al., 2018; Kong et al., 2018). For example, to generate the nuisance white matter signal, the white matter mask was defined anatomically and eroded three times. However, some have suggested that further erosion might be necessary to avoid the white matter signal being a hidden proxy for global signal (Power et al., 2017b). Indeed, in the GSP dataset, the white matter signal was quite correlated with the global signal with r = 0.77 ± 0.14 (mean ± std) across participants. Similarly, there is no consensus on selecting the DVARS threshold. We opted to select the DVARS threshold such that the number of censored frames due to DVARS was roughly the same as FD, which is the approach we considered in our previous studies (Kong et al., 2018), but might not be optimal (Afyouni and Nichols, 2018). Overall, there remains further room for exploring how the exact set of preprocessing hyperparameters might affect the final RSFC-behavior associations and behavioral prediction accuracies.

## 5. Conclusion

By applying the variance component model, we showed that GSR greatly increased the associations between behavior and RSFC in two large datasets of young, healthy adults. The results were replicated using kernel ridge regression and linear ridge regression (not shown). Given that GSR significantly reduced imaging-related artifacts, GSR-related improvements could not be simply explained by imaging artifacts being correlated with the behavioral measurements. The benefits of GSR on behavior-RSFC associations and prediction accuracies persisted even after ICA-FIX.

## Acknowledgement

This work was supported by Singapore MOE Tier 2 (MOE2014-T2-2-016), NUS Strategic Research (DPRT/944/09/14), NUS SOM Aspiration Fund (R185000271720), Singapore NMRC (CBRG/0088/2015), NUS YIA, the Singapore National Research Foundation (NRF) Fellowship (Class of 2017), and NIH K99AG054573. Our research also utilized resources provided by the Center for Functional Neuroimaging Technologies, P41EB015896 and instruments supported by 1S10RR023401, 1S10RR019307, and 1S10RR023043 from the Athinoula A. Martinos Center for Biomedical Imaging at the Massachusetts General Hospital. Our computational work for this article was partially performed on resources of the National Supercomputing Centre, Singapore (https://www.nscc.sg). Data were also provided by the Brain Genomics Superstruct Project of Harvard University and the Massachusetts General Hospital (Principal Investigators: Randy Buckner, Joshua Roffman, and Jordan Smoller), with support from the Center for Brain Science Neuroinformatics Research Group, the Athinoula A. Martinos Center for Biomedical Imaging, and the Center for Human Genetic Research. Twenty individual investigators at Harvard and MGH generously contributed data to the overall project. Data were also provided by the Human Connectome Project, WU-Minn Consortium (Principal Investigators: David Van Essen and Kamil Ugurbil; 1U54MH091657) funded by the 16 NIH Institutes and Centers that support the NIH Blueprint for Neuroscience Research; and by the McDonnell Center for Systems Neuroscience at Washington University.

## Appendix

In this appendix, we describe the mathematical details of kernel ridge regression. Let 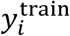 denote the score of a behavioral measure of training subject *i*. Let 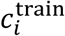 denote the vectorized RSFC (i.e., lower triangular entries of the RSFC matrix) of training subject *i*. The RSFC-based similarity between two training subjects *i* and *j* can be denoted as 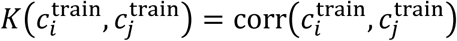, where corr(⋅) represents Pearson’s correlation.

Suppose there are *N*_1_ training subjects. Let ***K*** be an *N*_1_ × *N*_1_ matrix, where the *i*-th row and *j*-th column of ***K*** is *K*(*c*_*i*_, *c*_*j*_). Therefore, ***K*** is the similarity matrix among the *N*_1_ training subjects. In the training phase for a specific behavioral measure, the regression coefficient ***α*** is estimated from the training subjects via Eq. A1:

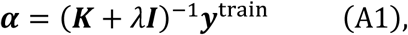

where ***y***^train^ is the *N*_1_ × 1 vector containing the behavioral scores of all training subjects. In this work, the regularization hyperparameter *λ* was selected by 20-fold cross-validation within the training set (i.e., inner-loop cross-validation).

Suppose the RSFC similarity between a test subject *s* and each training subject is consolidated into a 1 × *N*_1_ vector 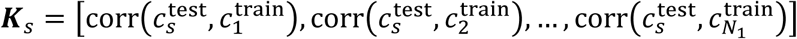, where 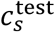 is the vectorized RSFC of the test subject. The predicted behavioral score of this test subject will be:

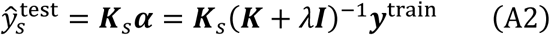

## Footnotes

1 Note that the cortical global signal is highly correlated with whole brain signal (r = 0.95 in the HCP dataset).

