## Supplemental materials for "Global Signal Regression Strengthens Association between Resting-State Functional Connectivity and Behavior"

### Supplementary Materials

| **Description** | **GSP field** |
| --- | --- |
| Flanker Accuracy | Flanker_S_CORRpc |
| Mental Rotation Accuracy | Average of MenRot_80_CORRpc, MenRot_120_CORRpc, and MentRot_160_CORRpc |
| Shipley Vocabulary | Shipley_Vocab_Raw |
| WAIS – Matrix Reasoning | Matrix_WAIS |
| Speilberger Trait Anxiety | STAI_tAnxiety |
| Speilberger State Anxiety | STAI_sAnxiety |
| Total Mood Disturbance | POMS_TotMdDisturb |
| Barratt Impulsivity | Barratt_tot |
| Mind Wandering Frequency | MindWandering_Freq |
| NEO – Neuroticism | NEO_N |
| NEO – Extraversion | NEO_E |
| NEO – Openness to Experience | NEO_O |
| NEO – Agreeableness | NEO_A |
| NEO – Conscientiousness | NEO_C |
| Novelty Seeking | TCI_Novelty |
| Reward Dependence | TCI_RewardDependance |
| Harm Avoidance | TCI_HarmAvoidance |
| Risk Raking | DOSPERT_taking |
| Risk Perception | DOSPERT_perception |
| Behavioral Activation – Drive | BISBAS_BAS_Drive |
| Behavioral Activation – Fun | BISBAS_BAS_Fun |
| Behavioral Activation – Reward | BISBAS_BAS_Reward |
| Behavioral Inhibition | BISBAS_BIS |

**Table S1.** Lookup table showing the original GSP variable names with the corresponding descriptive labels used in the manuscript.

| **Description** | **HCP field** |
| --- | --- |
| Visual Episodic Memory | PicSeq_Unadj |
| Cognitive Flexibility (DCCS) | CardSort_Unadj |
| Inhibition (Flanker Task) | Flanker_Unadj |
| Fluid Intelligence (PMAT) | PMAT24_A_CR |
| Vocabulary (Pronunciation) | ReadEng_Unadj |
| Vocabulary (Picture Matching) | PicVocab_Unadj |
| Processing Speed | ProcSpeed_Unadj |
| Delay Discounting | DDic_AUC_40K |
| Spatial Orientation | VSPLOT_TC |
| Sustained Attention – Sens. | SCPT_SEN |
| Sustained Attention – Spec. | SCPT_SPEC |
| Verbal Episodic Memory | IWRD_TOT |
| Working Memory (List Sorting) | ListSort_Unadj |
| Cognitive Status (MMSE) | MMSE_Score |
| Sleep Quality (PSQI) | PSQI_Score |
| Walking Endurance | Endurance_Unadj |
| Walking Speed | GaitSpeed_Unadj |
| Manual Dexterity | Dexterity_Unadj |
| Grip Strength | Strength_Unadj |
| Odor Identification | Odor_Unadj |
| Pain Interference Survey | PainInterf_Tscore |
| Taste Intensity | Taste_Unadj |
| Contrast Sensitivity | Mars_Final |
| Emotional Face Matching | Emotion_Task_Face_Acc |
| Arithmetic | Language_Task_Math_Avg_Difficulty_Level |
| Story Comprehension | Language_Task_Story_Avg_Difficulty_Level |
| Relational Processing | Relational_Task_Acc |
| Social Cognition – Random | Social_Task_Perc_Random |
| Social Cognition – Interaction | Social_Task_Perc_TOM |

**Table S2.** Lookup table showing the original HCP variable names with the corresponding descriptive labels used in the manuscript.

| **Description** | **HCP field** |
| --- | --- |
| Working Memory (N-back) | WM_Task_Acc |
| Agreeableness (NEO) | NEOFAC_A |
| Openness (NEO) | NEOFAC_O |
| Conscientiousness (NEO) | NEOFAC_C |
| Neuroticism (NEO) | NEOFAC_N |
| Extraversion (NEO) | NEOFAC_E |
| Emot. Recog. – Total | ER40_CR |
| Emot. Recog. – Angry | ER40ANG |
| Emot. Recog. – Fear | ER40FEAR |
| Emot. Recog. – Happy | ER40HAP |
| Emot. Recog. - Neutral | ER40NOE |
| Emot. Recog. – Sad | ER40SAD |
| Anger – Affect | AngAffect_Unadj |
| Anger – Hostility | AngHostil_Unadj |
| Anger – Aggression | AngAggr_Unadj |
| Fear – Affect | FearAffect_Unadj |
| Fear – Somatic Arousal | FearSomat_Unadj |
| Sadness | Sadness_Unadj |
| Life Satisfaction | LifeSatisf_Unadj |
| Meaning & Purpose | MeanPurp_Unadj |
| Positive Affect | PosAffect_Unadj |
| Friendship | Friendship_Unadj |
| Loneliness | Loneliness_Unadj |
| Perceived Hostility | PercHostil_Unadj |
| Perceived Rejection | PercReject_Unadj |
| Emotional Support | EmotSupp_Unadj |
| Instrument Support | InstruSupp_Unadj |
| Perceived Stress | PercStress_Unadj |
| Self-Efficacy | SelfEff_Unadj |

**Table S2 (cont.).** Lookup table showing the original HCP variable names with the corresponding descriptive labels used in the manuscript.

| **GSP behavior description** | **FD-behavior correlation** | **DVARS-behavior correlation** |
| --- | --- | --- |
| Harm Avoidance | -0.0899 | -0.0578 |
| NEO - Neuroticism | -0.0724 | -0.0519 |
| NEO - Agreeableness | -0.0702 | -0.0286 |
| Behavioral Inhibition | -0.0560 | -0.0792 |
| Flanker Accuracy | -0.0492 | -0.0662 |
| Speilberger Trait Anxiety | -0.0356 | -0.0016 |
| Total Mood Disturbance | -0.0302 | -0.0068 |
| NEO - Conscientiousness | -0.0278 | -0.0999 |
| Mind Wandering Frequency | -0.0268 | -0.0133 |
| Risk Perception | -0.0230 | -0.0896 |
| Speilberger State Anxiety | -0.0176 | -0.0237 |
| NEO - Extraversion | -0.0174 | 0.0218 |
| Behavioral Activation - Drive | -0.0137 | 0.0244 |
| Behavioral Activation - Reward | -0.0114 | 0.0108 |
| WAIS - Matrix Reasoning | -0.0113 | 0.0299 |
| Reward Dependence | -0.0105 | -0.0520 |
| Barratt Impulsivity | -0.0035 | 0.0516 |
| NEO - Openness to Experience | 0.0063 | -0.0307 |
| Behavioral Activation - Fun | 0.0226 | 0.0599 |
| Shipley Vocabulary | 0.0342 | 0.0646 |
| Novelty Seeking | 0.0383 | 0.0559 |
| Mental Rotation Accuracy | 0.0433 | 0.0429 |
| Risk Taking | 0.0659 | 0.0646 |

**Table S3.** Pearson’s correlation between motion (FD and DVARS) and the behavioral scores in the GSP dataset.

| **HCP behavior description** | **FD-behavior correlation** | **DVARS-behavior correlation** |
| --- | --- | --- |
| Walking Endurance | -0.2973 | 0.0685 |
| Working Memory (N-back) | -0.2361 | 0.0151 |
| Reading (Pronounciation) | -0.2143 | 0.0390 |
| Fluid Intelligence (PMAT) | -0.1931 | 0.0399 |
| Vocabulary (Picture Matching) | -0.1864 | 0.0577 |
| Relational Processing | -0.1834 | -0.0403 |
| Spatial Orientation | -0.1616 | 0.1028 |
| Delay Discounting | -0.1521 | 0.0454 |
| Cognitive Flexibility (DCCS) | -0.1461 | 0.0085 |
| Working Memory (List Sorting) | -0.1372 | 0.0462 |
| Openness (NEO) | -0.1283 | 0.0788 |
| Manual Dexterity | -0.1235 | -0.0880 |
| Story Comprehension | -0.1122 | 0.0824 |
| Agreeableness (NEO) | -0.1071 | -0.1356 |
| Emot. Recog. - Total | -0.1044 | -0.0721 |
| Social Cognition - Interaction | -0.1036 | 0.0364 |
| Visual Episodic Memory | -0.1034 | -0.0977 |
| Inhibition (Flanker task) | -0.0990 | 0.0015 |
| Contrast Sensitivity | -0.0955 | 0.1298 |
| Processing Speed | -0.0875 | 0.0604 |
| Verbal Episodic Memory | -0.0865 | -0.0387 |
| Arithmetic | -0.0831 | 0.0020 |
| Walking Speed | -0.0798 | -0.0949 |
| Emot. Recog. - Sad | -0.0700 | -0.0120 |
| Emot. Recog. - Angry | -0.0614 | -0.0352 |
| Emotional Face Matching | -0.0584 | -0.0301 |
| Emot. Recog. - Fear | -0.0568 | -0.0752 |
| Sustained Attention - Spec. | -0.0457 | -0.0837 |
| Life Satisfication | -0.0455 | -0.0146 |

**Table S4.** Pearson’s correlation between motion (FD and DVARS) and the behavioral scores in the HCP dataset.

| **HCP behavior description** | **FD-behavior correlation** | **DVARS-behavior correlation** |
| --- | --- | --- |
| Emot. Recog. - Happy | -0.0421 | 0.0119 |
| Emot. Recog. - Neutral | -0.0380 | -0.0400 |
| Loneliness | -0.0362 | 0.0026 |
| Cognitive Status (MMSE) | -0.0288 | -0.0572 |
| Odor Identificaiton | -0.0228 | -0.0958 |
| Emotional Support | -0.0213 | -0.0260 |
| Instrument Support | -0.0207 | 0.0058 |
| Sadness | -0.0204 | -0.0315 |
| Extraversion (NEO) | -0.0085 | 0.0332 |
| Sustained Attention - Sens. | -0.0062 | 0.0567 |
| Fear - Affect | -0.0056 | -0.0557 |
| Conscientiousness (NEO) | -0.0046 | -0.0689 |
| Neuroticism (NEO) | 0.0098 | -0.0382 |
| Perceived Stress | 0.0231 | -0.0421 |
| Grip Strength | 0.0241 | 0.3659 |
| Pain Interference Survey | 0.0256 | 0.0273 |
| Anger - Affect | 0.0258 | 0.0065 |
| Anger - Hostility | 0.0354 | 0.0320 |
| Friendship | 0.0385 | 0.0411 |
| Self-Efficacy | 0.0424 | 0.0945 |
| Perceived Hostility | 0.0427 | 0.0690 |
| Social Cognition - Random | 0.0478 | 0.0093 |
| Perceived Rejection | 0.0482 | -0.0069 |
| Positive Affect | 0.0555 | 0.0456 |
| Fear - Somatic Arousal | 0.0558 | 0.0093 |
| Taste Intensity | 0.0737 | -0.0744 |
| Anger - Aggression | 0.0760 | 0.1236 |
| Meaning & Purpose | 0.0761 | 0.0167 |
| Sleep Quality (PSQI) | 0.1299 | 0.0115 |

**Table S4 (cont.).** Pearson’s correlation between motion (FD and DVARS) and the behavioral scores in the HCP dataset.

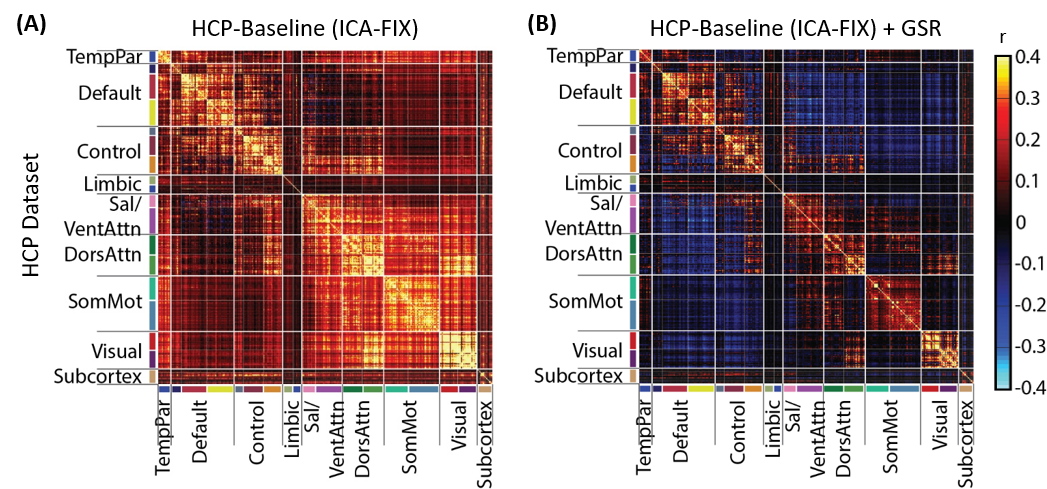

**Figure S1. GSR results in a negative “shift” in RSFC in the HCP dataset.** (A) RSFC matrix among the 419 ROIs using baseline processing without GSR. (B) RSFC matrix among the 419 ROIs using baseline processing with GSR. For visualization, the 419 ROIs are ordered according to the 17 networks in Figure 1A and subcortical structures listed in Figure 1B. These 17 networks are in turn divided into eight groups (TempPar, Default, Control, Limbic, Salience/Ventral Attention, Dorsal Attention, Somatomotor and Visual), roughly corresponding to major networks discussed in the literature. These eight groups and subcortical structures are separated by thick while lines. Consistent with the literature, GSR introduces a negative shift in the RSFC.

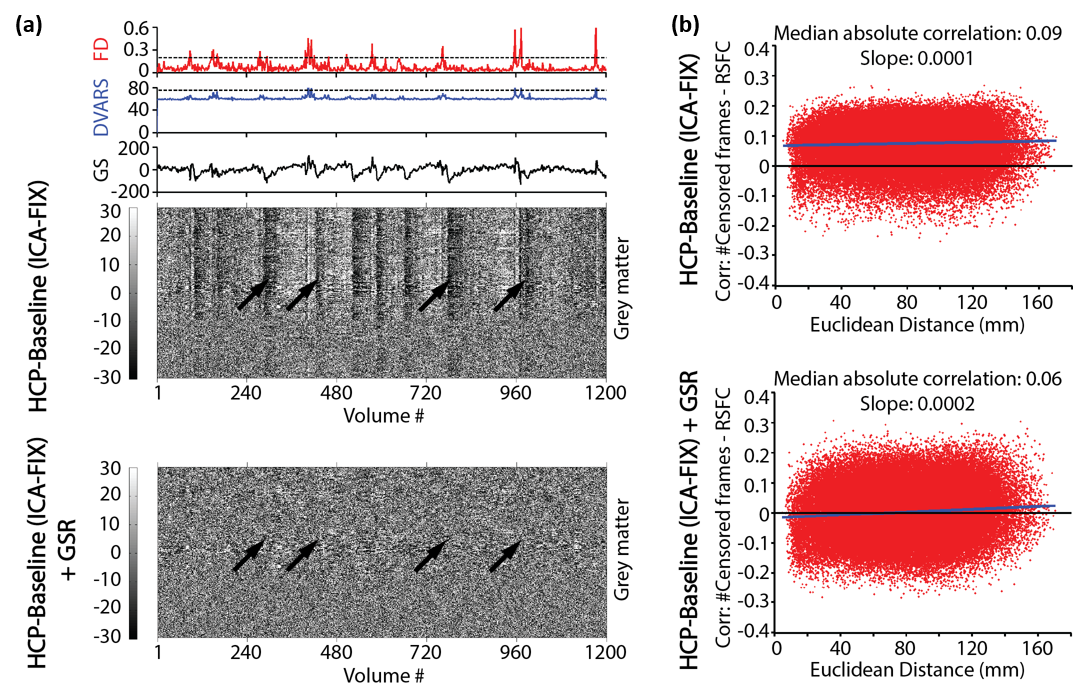

**Figure S2. GSR reduces imaging artifacts, while exacerbating distance-dependent FC biases in the HCP dataset.** (A) QC plot of a representative mid-motion HCP subject. FD (red), DVARS (blue), and GS (black) are shown in the top three panels. The horizontal lines in the first two panels indicate the thresholds used in the censoring step. The bottom two panels are signal intensity of gray matter voxels in two preprocessing pipelines: HCP-Baseline (upper) and HCP-Baseline+GSR (lower). Without GSR, motion-related global artifacts are visible in the gray matter timeseries, while GSR removes the global signal changes (black arrows). (B) Correlations between QC (number of censored frames) and RSFC are shown for two preprocessing pipelines: HCP-Baseline (upper) and HCP-Baseline+GSR (lower). Each red dot indicates one ROI pair. The x-axes are the Euclidean distances among pairs of ROIs. The functional connectivity between each pair of ROIs are correlated with the number of censored frames across all subjects and shown on the y-axes. The blue lines corresponded to linear fits of the red dots.

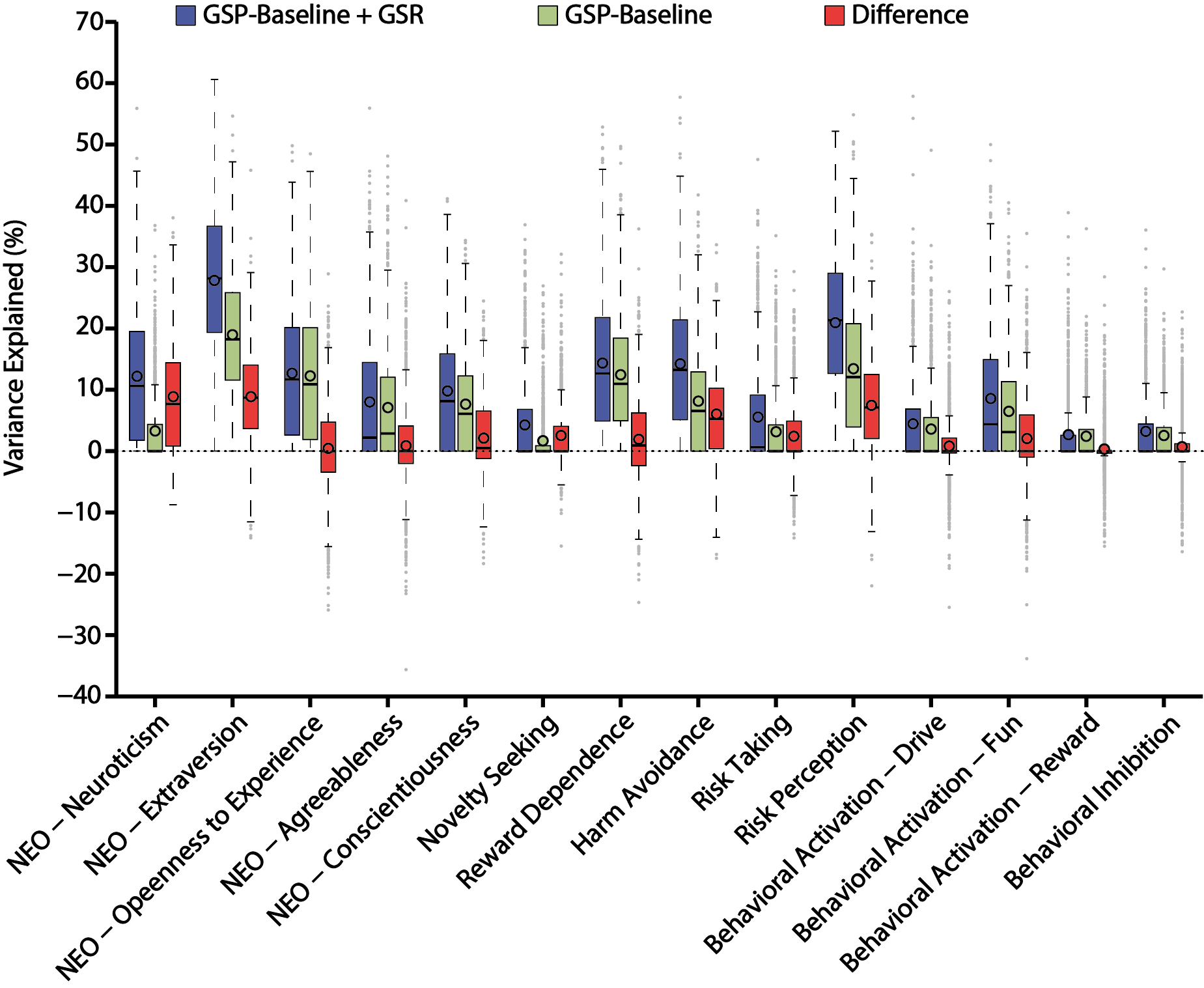

**Figure S3.** **GSR improves behavioral variance explained by RSFC in the GSP dataset**. Behavioral variance explained by RSFC for 14 remaining measures of personality and associated biosocial factors. For each behavioral measure, the explained variance by the GSP-Baseline+GSR pipeline, GSP-Baseline pipeline, and difference between the two pipelines are shown in blue, green and red respectively. The “boxes” show the median and interquartile range (IQR) of explained variance across all jackknife samples. The whisker length is 1.5 IQR. Black circles indicate mean. Outliers are shown by grey dots.

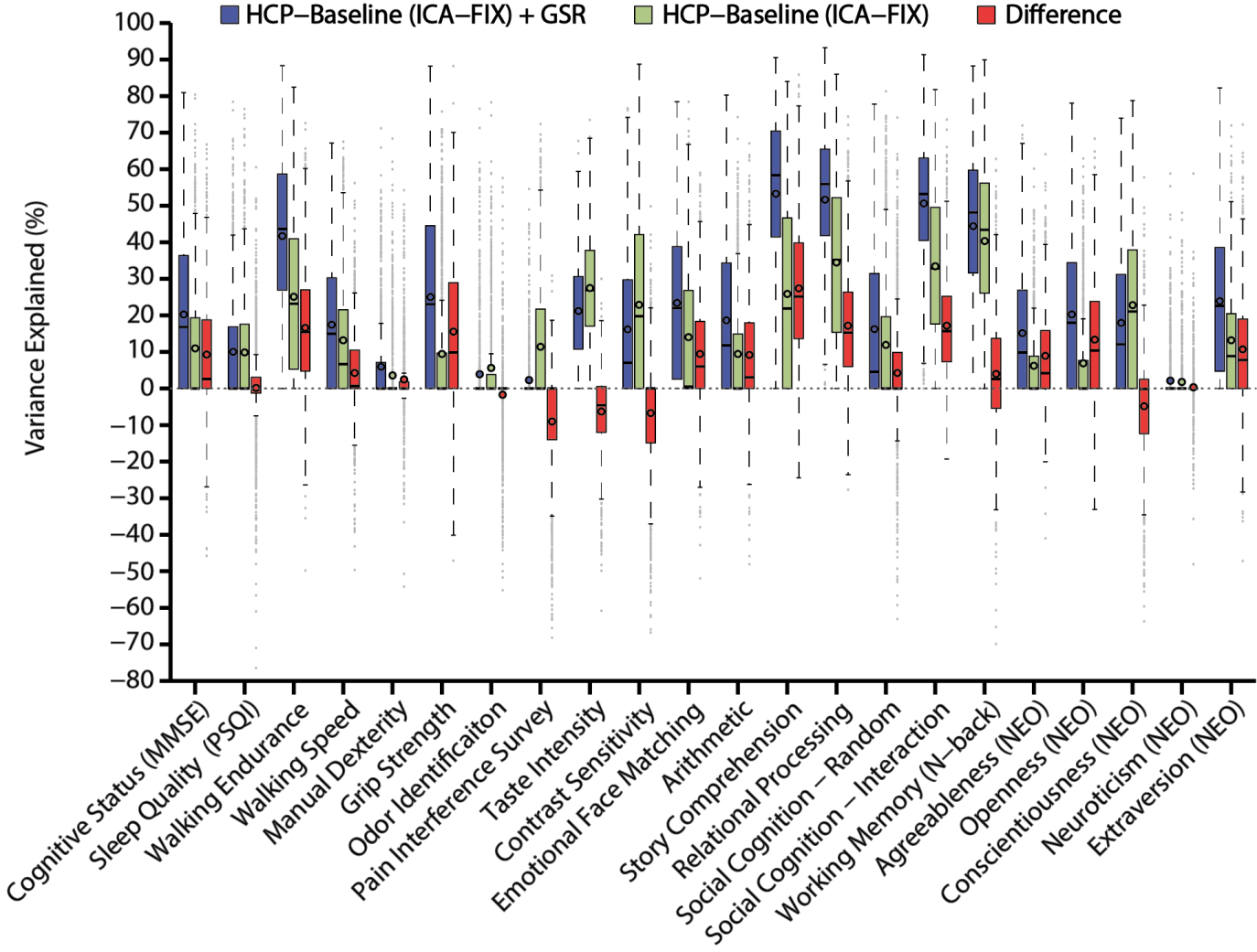

**Figure S4. GSR improves behavioral variance explained by RSFC in the HCP dataset**. Behavioral variance explained by RSFC for 22 measures including alertness, motor, sensory, personality, and in-scanner measures. For each behavioral measure, the explained variance by the HCP-Baseline+GSR pipeline, HCP-Baseline pipeline, and difference between the two pipelines are shown in blue, green and red respectively. The “boxes” show the median and interquartile range (IQR) of explained variance across all jackknife samples. The whisker length is 1.5 IQR. Black circles indicate mean. Outliers are shown by grey dots.

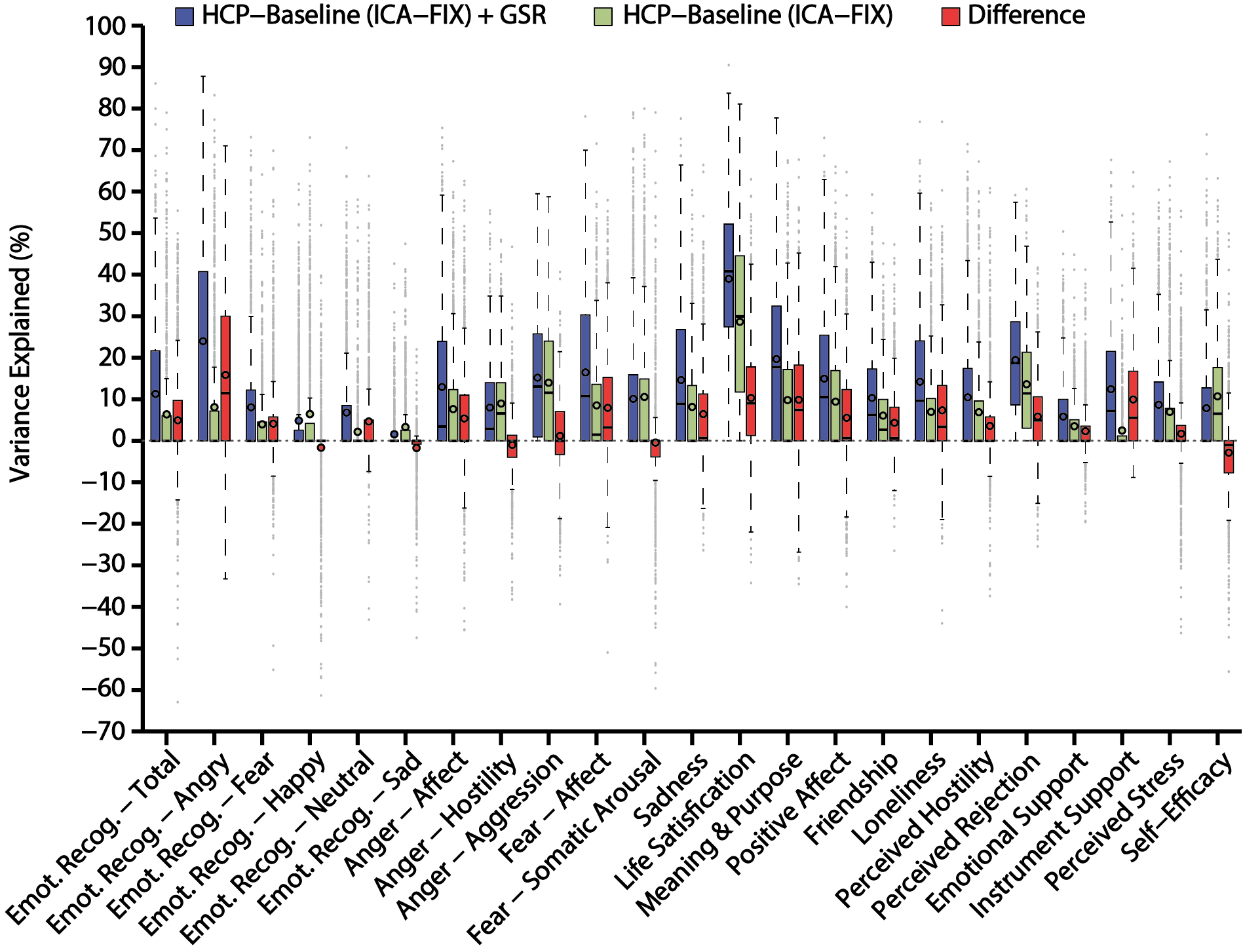

**Figure S5. GSR improves behavioral variance explained by RSFC in the HCP dataset**. Behavioral variance explained by RSFC for 23 social-emotional measures. For each behavioral measure, the explained variance by the HCP-Baseline+GSR pipeline, HCP-Baseline pipeline, and difference between the two pipelines are shown in blue, green and red respectively. The “boxes” show the median and interquartile range (IQR) of explained variance across all jackknife samples. The whisker length is 1.5 IQR. Black circles indicate mean.

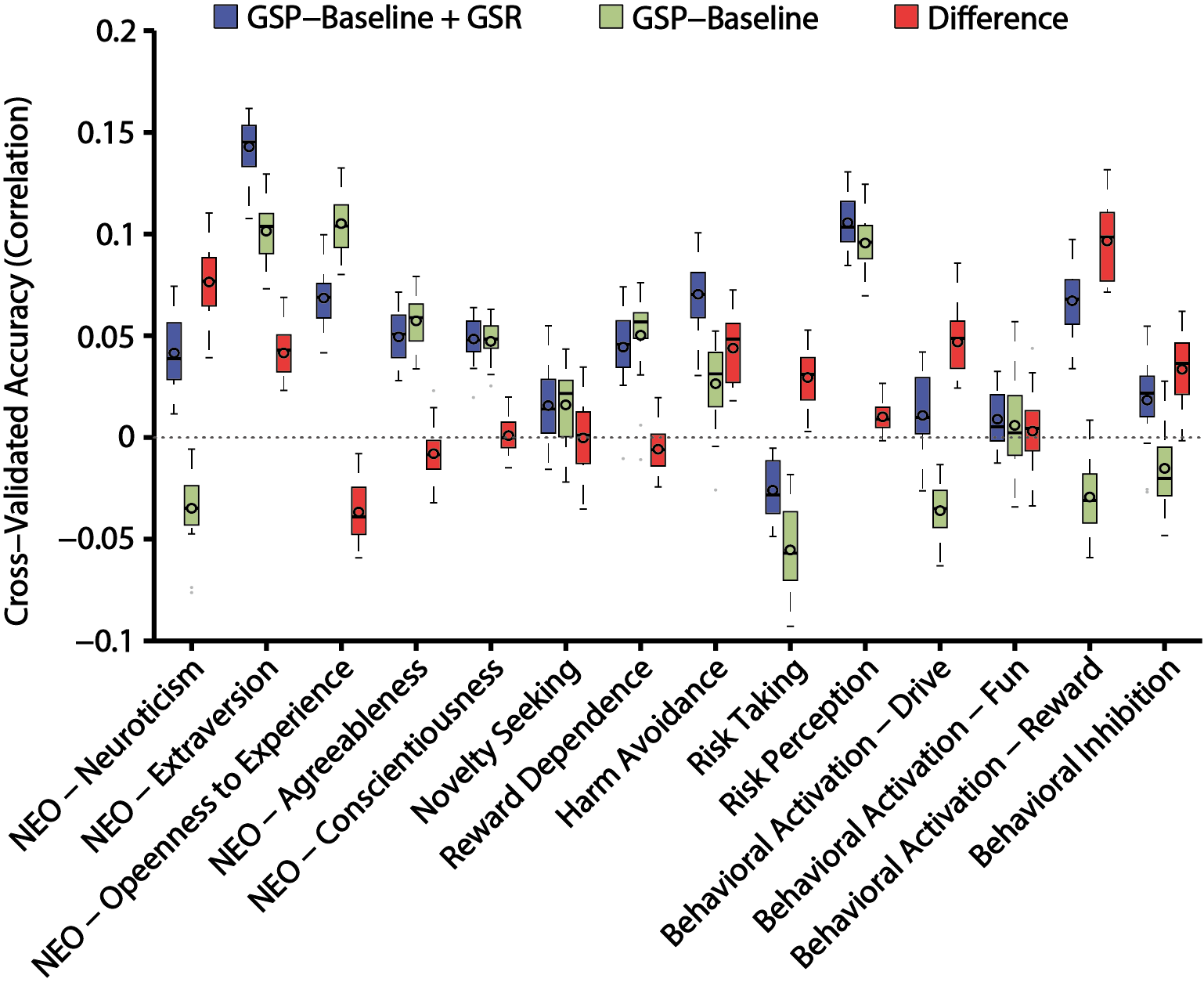

**Figure S6.** **GSR improves RSFC-based behavioral prediction accuracies in the GSP dataset using kernel ridge regression**. Test accuracies for 14 remaining measures of personality and associated biosocial factors. For each behavioral measure, the accuracies with the GSP-Baseline+GSR pipeline, GSP-Baseline pipeline, and difference between the two pipelines are shown in blue, green and red respectively. The “boxes” show the median and interquartile range (IQR) of test accuracies across 20 random cross-validation splits. The whisker length is 1.5 IQR. Black circles indicate mean. Outliers are shown by grey dots.

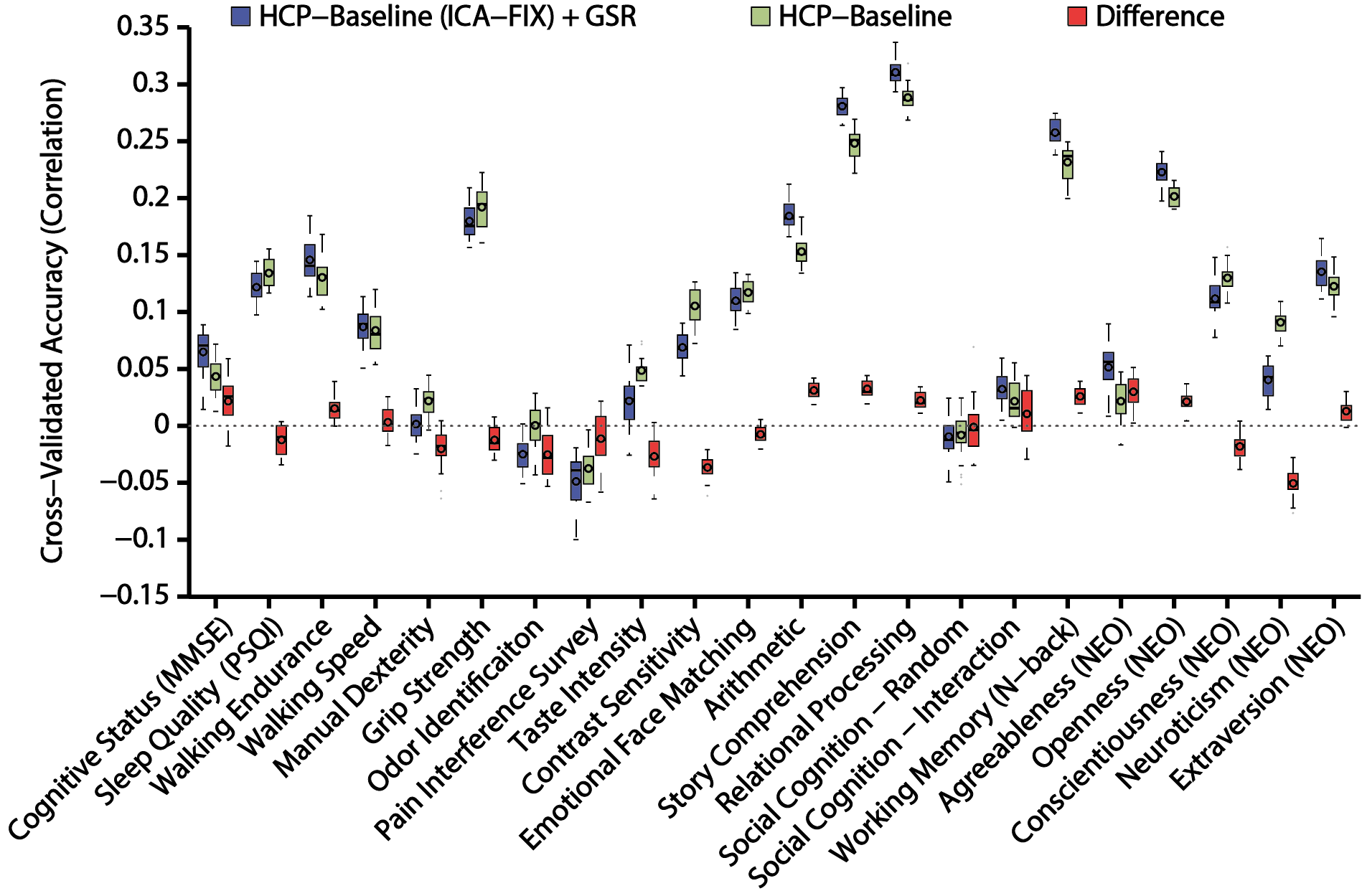

**Figure S7. GSR improves RSFC-based behavioral prediction accuracies in the HCP dataset using kernel ridge regression**. Test accuracies for 22 measures including alertness, motor, sensory, personality, and in-scanner measures. For each behavioral measure, the accuracies with the HCP-Baseline+GSR pipeline, HCP-Baseline pipeline, and difference between the two pipelines are shown in blue, green and red respectively. The “boxes” show the median and interquartile range (IQR) of test accuracies across 20 random cross-validation splits. The whisker length is 1.5 IQR. Black circles indicate mean. Outliers are shown by grey dots.

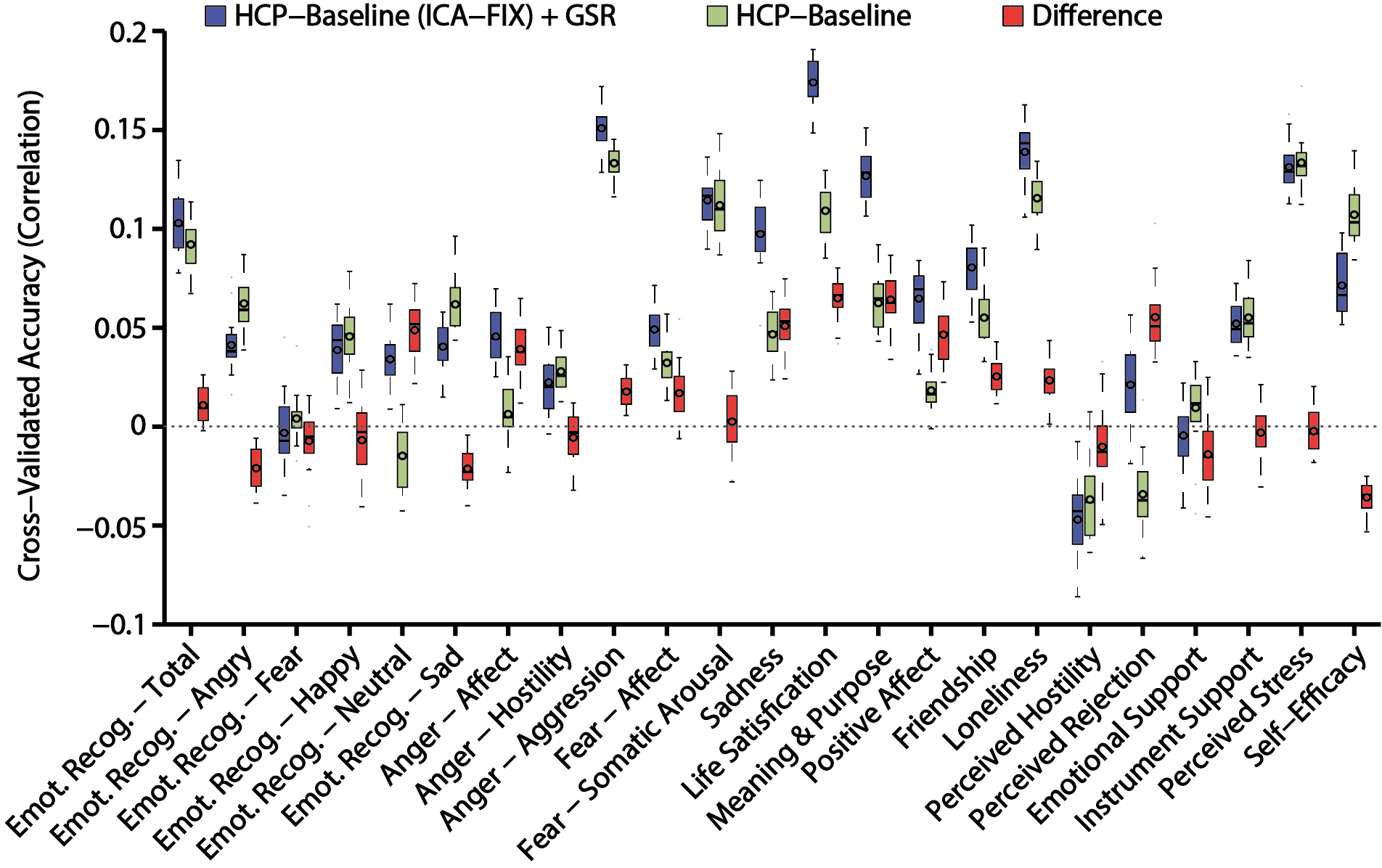

**Figure S8. GSR improves RSFC-based behavioral prediction accuracies in the HCP dataset using kernel ridge regression**. Test accuracies for 23 social-emotional measures. For each behavioral measure, the accuracies with the HCP-Baseline+GSR pipeline, HCP-Baseline pipeline, and difference between the two pipelines are shown in blue, green and red respectively. The “boxes” show the median and interquartile range (IQR) of test accuracies across 20 random cross-validation splits. The whisker length is 1.5 IQR. Black circles indicate mean.

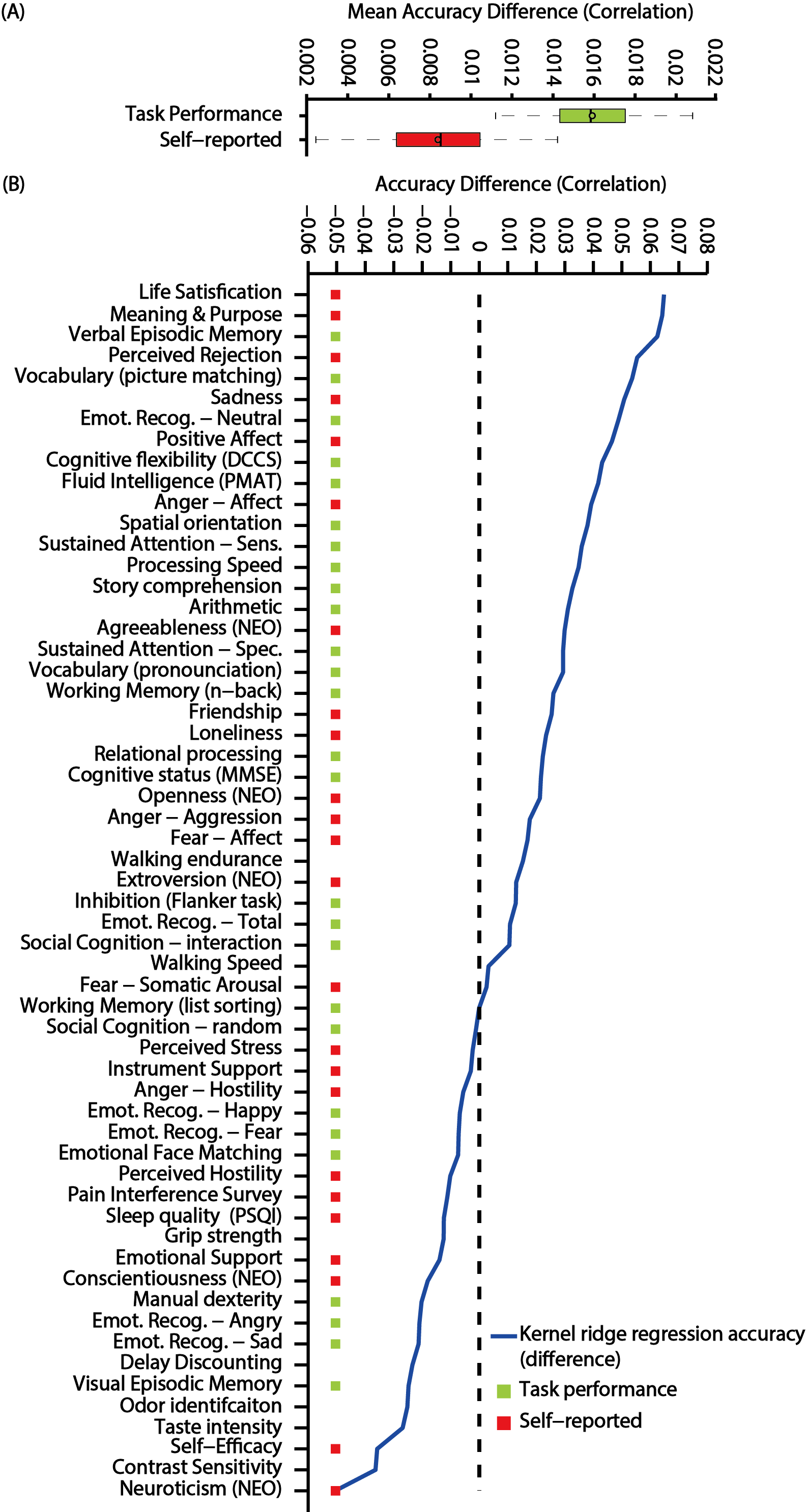

**Figure S9. GSR preferentially improves behavioral prediction accuracies for task-performance measures compared with self-reported measures using kernel ridge regression.** (A) Improvement in test accuracies averaged across all HCP task-performance measures (green) and across all self-reported measures (red). The “boxes” show the median and interquartile range (IQR) of test accuracies across all random data splits. The whisker length is 1.5 IQR. Black circles indicate mean. (B) Behavioral measures are ordered based on improvement in test accuracies, i.e., baseline+GSR minus baseline (blue line). Behavioral measures marked with green color are considered task-performance measures. Behavioral measures marked with red color are considered self-reported measures.
